# Peripheral axonal ensheathment is regulated by Ral GTPase and the exocyst complex

**DOI:** 10.1101/537969

**Authors:** Joana F. Silva-Rodrigues, Cátia F. Patrício-Rodrigues, Vicente de Sousa-Xavier, Pedro M. Augusto, Ana C. Fernandes, Ana R. Farinho, Rita O. Teodoro

## Abstract

Axon ensheathment is fundamental for fast impulse conduction and the normal physiological functioning of the nervous system. Defects in axonal insulation lead to debilitating conditions, but despite its importance, the molecular players responsible are poorly defined. Here, we identify Ral GTPase as a key player in axon ensheathment in *Drosophila* larval peripheral nerves. We demonstrate through genetic analysis that Ral action through the exocyst complex is sufficient and necessary in wrapping glial cells to regulate their growth and development. We suggest that the Ral-exocyst pathway controls the targeting of secretory vesicles for membrane growth or for the secretion of a wrapping glia-specific factor that itself regulates growth. In summary, our findings provide a new molecular understanding of the process by which axons are ensheathed *in vivo*, a process critical for normal neuronal function.

## Introduction

The peripheral nervous system (PNS) is responsible for the innervation of the body musculature and organs, and to relay signals from the periphery to the brain. Timely delivery of information across body regions is critical for adequate neuronal function and behavior. Fast and efficient conduction of the nerve impulse requires appropriate axonal insulation, known to be mediated by glial wrapping of peripheral axons. In vertebrates, this function is assured by myelinating and non-myelinating Schwann cells, which are responsible for ensheathing and supporting the axons within peripheral nerves. Defects in axonal insulation lead to debilitating conditions, including multiple sclerosis (Saab & Nave 2017; Osso & Chan 2017; Chang et al. 2016; Ben A Barres 2008). In invertebrates, like *Drosophila*, PNS axons are similarly enwrapped by glial cells, but instead of producing myelin, they form a myelin-like sheath by producing large amounts of membrane (Schirmeier et al. 2015; Banerjee & Bhat 2008; Silies et al. 2007; Rodrigues et al. 2010). Defects in wrapping glia give rise to increased spike propagation times (Ghosh et al. 2013). How glial and neuronal development is coordinated and how these distinct cell types develop and interact to ensure correct nervous system function and normal behavior is far from being understood.

*Drosophila* larval peripheral nerves are composed of descending motor and ascending sensory neuron axons that are surrounded by three distinct glial layers, which are then enclosed by an acellular extracellular matrix (ECM) layer - called lamella. The outermost glial layer is made of perineurial glia, whose function remains elusive. Following this layer are subperineurial glia, which form auto-septate junctions (SJ) and are responsible for the formation of the blood nerve barrier (BNB), thought to prevent the potassium-rich haemolymph to be in contact with the axons. Below the subperineurial glia, is the innermost glial layer formed by wrapping glia, the glial subtype that assures ensheathment and insulation of axons. Unlike vertebrate Schwann cells, wrapping glia do not produce myelin, but grow tremendously during development producing large amounts of membrane that assure that axons are either wrapped individually or in small bundles (Stork et al. 2008; Silies et al. 2007; Matzat et al. 2015; Hilchen et al. 2013). The formation of glial sheaths starts during embryonic development and proceeds throughout larval stages, where the larvae increase in size ∼100 times. During larval development, wrapping glia and subperineurial glia show hypertrophic growth without proliferation, while perineurial glia become proliferative and divide (Hilchen et al. 2013). This means that all wrapping glia and subperineurial glia present at the end of larval stages are generated during embryogenesis and their membranes have to massively expand to accommodate larval growth (Matzat et al. 2015; Hilchen et al. 2013; Hilchen et al. 2008). The mechanism(s) by which these growth processes are regulated remain largely mysterious, with very few genes identified as required for the development of these distinct glial subtypes.

In the *Drosophila* PNS, a few factors have been reported to play a role in wrapping glia development (Matzat et al. 2015; Ghosh et al. 2013; Schmidt et al. 2012; Xie & Auld 2011; Leiserson et al. 2000; Lavery et al. 2007), but while these studies revealed some of the signaling pathways, how this enormous membrane-remodeling event is specifically regulated is poorly understood. Given the nature of this process, genes involved in membrane addition are likely to be required. A pathway that has recurrently been reported to orchestrate the polarized targeting of vesicles and to regulate membrane growth events is the exocyst complex, via its interaction with the small GTPase Ral (Lee & Schwarz 2016; Teodoro et al. 2013; Armenti et al. 2014; Balasubramanian et al. 2010; Patrício-Rodrigues & Teodoro 2018). The exocyst is an octameric protein complex conserved from yeast to human that is an effector to many GTPases, including Ral. By being able to receive regulatory information from different pathways, the exocyst can serve as a hub to precisely regulate where and when vesicles fuse with the membrane (Mei & Guo 2018; Picco et al. 2017; Martin-Urdiroz et al. 2016). Interaction with Ral, a small GTPase from the Ras superfamily, induces the assembly and activation of this complex (Moskalenko et al. 2001). Interestingly, one of the mammalian Ral isoforms, RalA, is present in oligodendrocytes (OL) where it colocalizes to a relatively high degree with Proteolipid protein, a myelin constituent, leading to the suggestion that RalA can be involved in myelin membrane biogenesis (Anitei et al. 2009), but this remains to be tested. Likewise, the exocyst subunits Sec6 and Sec8 are present in OLs and in myelin, and Sec8 has been shown to participate in OL differentiation - a process characterized by rapid process extension, membrane expansion and polarization (Anitei 2006). Additionally, through an interaction with Dlg1, Sec8 has also been shown to contribute to myelin formation in culture (Bolis et al. 2009). Together, these data hint at a function for Ral and for the exocyst in the process of axonal wrapping, but whether they are required for myelin formation or axonal ensheathing, or whether they are in the same pathway, is not known.

Here we show that Ral GTPase and the exocyst are critical regulators of nerve bundle development in the *Drosophila* PNS. We demonstrate that *ral* mutants have severely underdeveloped wrapping glia but have morphologically normal subperineurial and perineurial glia, implying a specific role for Ral GTPase in wrapping glia and not in general growth. We show that wrapping glia defects result from a failure to grow rather than a failure to maintain membrane integrity. In addition, we demonstrate that these mutants have thicker ISNs, often with defasciculated axons and signs of a disrupted BNB. These defects in the mutants result in abnormal locomotion, possibly due to altered axonal insulation. Genetic analyses show that Ral is both sufficient and necessary in wrapping glia to regulate its growth and development. Furthermore, we show that Ral GTPase functions via the exocyst complex in the regulation of these developmental processes. Given the known roles of this pathway, we propose that the Ral/exocyst pathway regulates axonal wrapping directly through the control of the targeting of secretory vesicles for membrane addition or, alternatively, by controlling the secretion of a wrapping glia-specific factor, that would itself regulate growth. In summary, our findings establish Ral GTPase and the exocyst as novel players required for wrapping glia development, and axonal ensheathment *in vivo*, a process critical for normal impulse conduction.

## Results

### ral mutants have abnormal nerve bundles

Postembryonic PNS development requires substantial growth to accommodate the increase in larval size. Specifically, PNS axons and their surrounding glial cells must grow coordinately and with the rest of the body to assure the proper formation of nerve bundles (Figure 1A). Despite important for nerve impulse propagation the molecular pathways that regulate nerve bundle development are poorly understood. Knowing that Ral GTPase is enriched in myelin-producing oligodendrocytes, together with its known roles in the regulation of growth and proliferation in other cellular contexts, led us to hypothesize that Ral could participate in *Drosophila* larvae PNS development. By analyzing two distinct Ral GTPase mutants, *ral*^*G0501*^ and *ral*^*EE1*^, we observed that the 3^rd^ instar larval ISN bundles were abnormally thick in these mutants (Figure 1B-C). This phenotype is fully penetrant with every larvae analyzed in both *ral* mutants showing this defect in several larval segments. *ral*^*G0501*^ is a protein null mutant (Teodoro et al. 2013) and *ral*^*EE1*^ harbors a point mutation that results in an amino acid substitution, Ser154Leu, within a conserved amino acid sequence predicted to be required for nucleotide binding (Eun et al. 2006). We used anti-HRP to label the nerve outline and quantified the width of the ISN at muscle 4 (m4) region (see methods). Our data shows that wild-type (WT) larvae have an ISN that is 5.21±0.08 μm, which is significantly different from *ral*^*G0501*^ and *ral*^*EE1*^ where the ISN is thicker being 8.01±0.26μm and 7.82±0.13μm, respectively (Figure 1C-D). To minimize possible biases in the place of measurement of the nerve width, we calculated the nerve area corresponding to 80μm in length on the same region of ISN, in WT and *ral* mutants. In agreement to the increase in nerve width, we found that *ral* mutants have increased nerve area compared to WT (Figure 1E).

**Figure 1.**
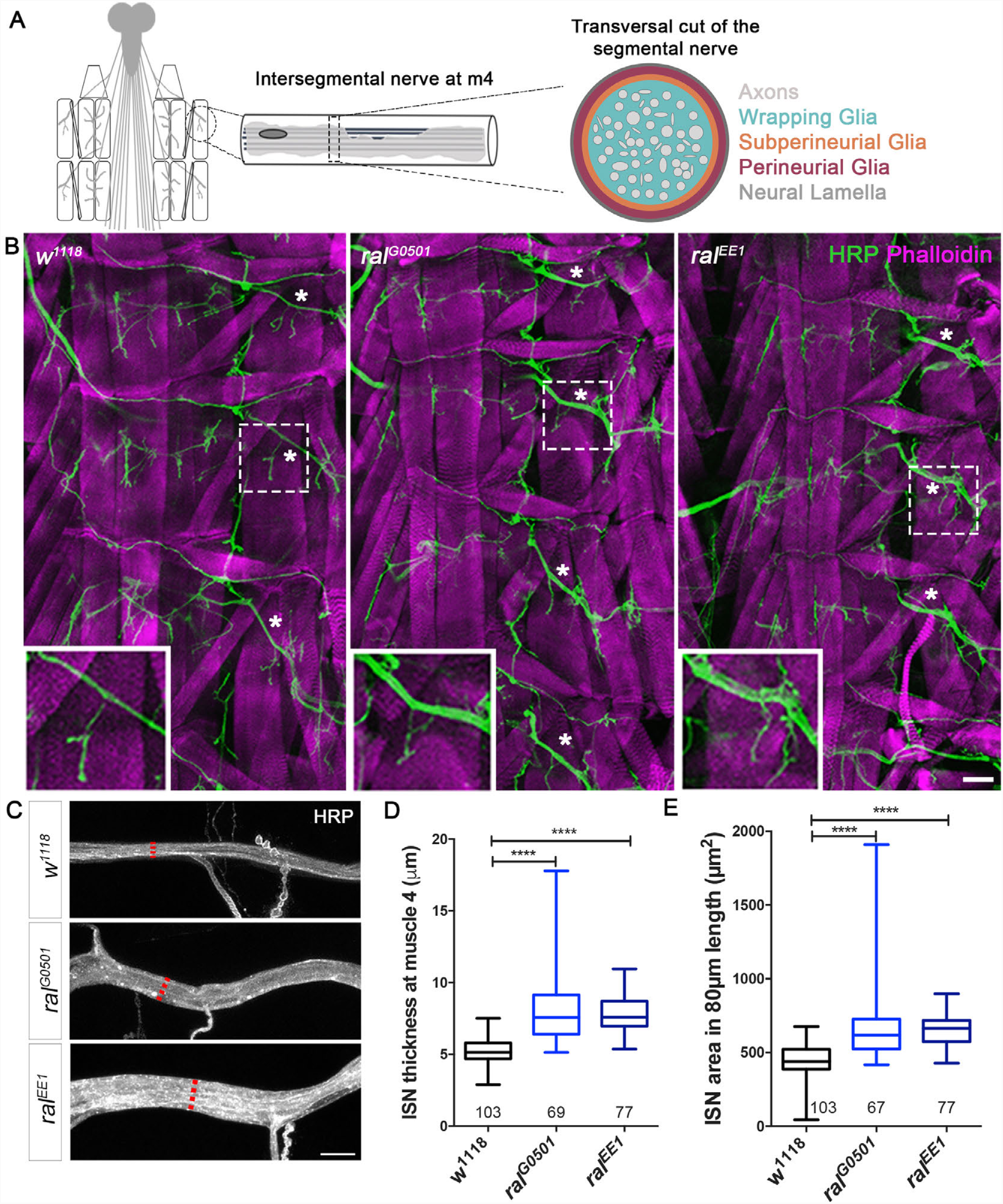
Ral GTPase mutant larvae have thicker intersegmental nerve (ISN) bundles than wild-type. **(A)** Schematic representation of (left to right): 1) a dissected fillet of a 3^rd^-instar larvae, with brain, nerves, muscles and neuromuscular junction represented, 2) longitudinal diagram of ISN, highlighting axons and glia, 3) cross-section of ISN representing all its components, colour-coded legend. **(B)** Representative example of wild-type (WT, *w*^*1118*^) and *ral* mutants (*ral*^*G0501*^ and *ral*^*EE1*^) 3^rd^-instar larval body walls showing muscles labelled with Phalloidin (magenta) and neuronal tissue labelled with HRP (green), emphasising the ISN (boxed area and zoom below). Arrows point towards the ISNs. Scale bar: 50μm. **(C)** Representative examples of ISN at muscle 4 (m4) exit point of wild-type (*w*^*1118*^), *ral*^*G0501*^ and *ral*^*EE1*^ mutant larvae showing increased nerve width of the mutants compared to wild-type. Dashed line shows the approximate place where thickness measurements are performed. Scale bar: 10μm. **(D)** Quantifications of the width of the ISN at m4 exit point. **(E)** Quantification of nerve area, per 80μm, confirmed that *ral*^*G0501*^ and *ral*^*EE1*^ have increased nerve area. In **(D)** and **(E)** numbers on x-axis represent the total number of ISNs quantified from segments A2-A4. Non-parametric ANOVA, Kruskal-Wallis with Dunn’s multiple comparisons test was used. ****p<0.0001.

### Ral GTPase is required for the development of wrapping glia

Alterations in nerve morphology can have several origins, including defects in glial development or in axonal fasciculation. In fact, it has been shown that defects in glia development are many times the underlying cause of nerve thickenings (Ghosh et al. 2013; Petley-Ragan et al. 2016; Schmidt et al. 2012; Leiserson et al. 2000; Sepp et al. 2001). To learn about the cellular origin of the observed defects we analyzed the morphology of each of these cell types (neurons and glia) by labeling each of them with CD4-GFP, expressed under the control of cell type-specific Gal4 drivers. Neurons were visualized using nSyb-Gal4, which labels all neurons (Figure 2A); for the different glia, we used Bsg-Gal4, Moody-Gal4 and Nrv2-Gal4 to label perineurial, subperineurial and wrapping glia, respectively (Figure 2B-D). Finally, the neural lamella was examined by analyzing the protein-trap Viking-GFP (Vkg-GFP) (Bainton 2014), a collagen-IV protein, and a major ECM protein (Figure 2E). We imaged the ISN from segments A2-A4 (Figure 2). Imaging of CD4-GFP and HRP allowed visualization of the nerve simultaneously with each of the cell types, and assess whether the morphology was altered in the mutants. Labeling of neurons with CD4-GFP revealed a certain degree of defasciculation in the mutants compared to control (Figure 2A). However, careful observation of the body wall innervation shows that there are no ectopic or de-innervated muscles (Figure 1B), implicating that neurons are present in the correct number and targeting to the correct location. Staining of axons with Futsch, an axonal microtubule-associated protein, was consistent with these observations revealing that axonal tracts were often defasciculated in *ral* mutants (Supplementary Figure 1), but in contrast with the increased thickness of ISN that was present in all mutant larvae, the defects in axonal fasciculation were more variable and less obvious, suggesting that defasciculation *per se* is unlikely to be the only cause for the alteration in nerve morphology.

**Figure 2.**
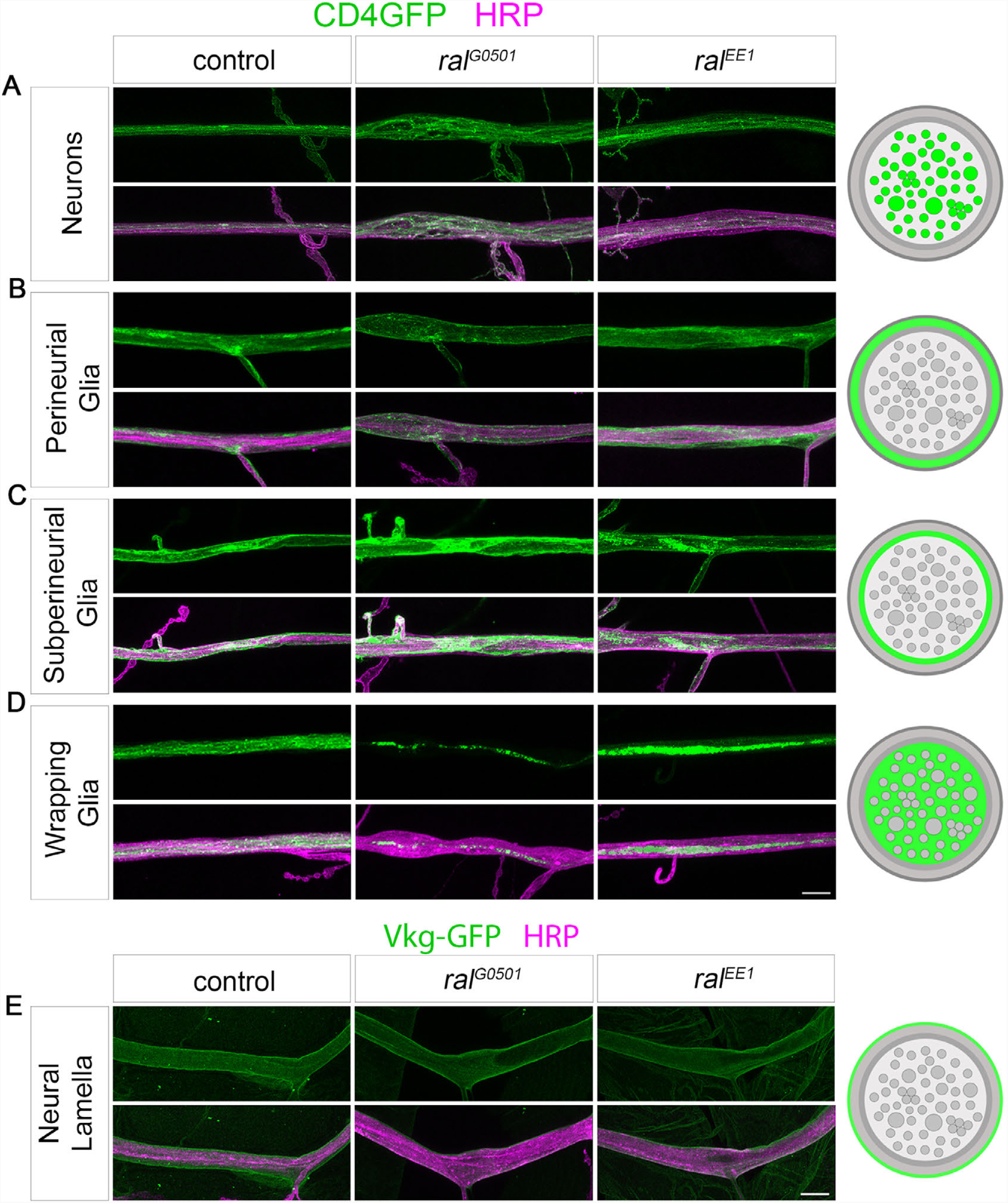
*ral* mutants have defective wrapping glia. **(A-D)** Analysis of the morphology of **(A)** Neurons, **(B)** Perineurial Glia, **(C)** Subperineurial Glia and **(D)** Wrapping Glia at ISN at m4 exit point. UAS-CD4GFP was used as a membrane marker, and cell-type specific Gal4 drivers (nSyb-Gal4>neurons, Bsg-Gal4>PG, moody-Gal4>SPG and Nrv2-Gal4>WG) were used. Membranes of each cell type were visualized by CD4-GFP (green) and neuronal tissue with HRP (magenta). For neural lamella **(E)** the protein trap Vkg-GFP (green) was used. The labelling was performed in *w*^*1118*^, *ral*^*G0501*^ and *ral*^*EE1*^ background. Wrapping glia are strikingly underdeveloped and fail to wrap the axons within the bundle in *ral*^*G0501*^ and *ral*^*EE1*^ larvae. Scale bar:10 µm. Schematic of the labeled cell type (green) on the right side, the non-labeled layers are represented in grey.

By extending this approach to glia, we observed that the overall morphology of perineurial and subperineurial glia in *ral* mutants is identical to controls (Figure 2B-C). Normally, perineurial glia wrap around the entire bundle, and this configuration is unchanged in *ral* mutants (Figure 2B). Likewise, subperineurial glia extend membrane processes that cover the entire length of the nerve bundle of the larvae, and this morphology is also unchanged in *ral* mutants (Figure 2C). In contrast with perineurial and subperineurial glia, the innermost glial layer, composed by wrapping glia, is severely underdeveloped in the mutants (Figure 2D). While in controls wrapping glia surround and completely wrap the axons, in both *ral* mutants the membranes of wrapping glia fail to grow and look like thin membrane extensions that fail to enclose the axons, a situation radically different from what is observed in controls (Figure 2A,D). Given that wrapping glia do not divide during larval development, it is likely that this phenotype results from a failure in cell growth and in membrane addition. As subperineurial glia also grow and do not divide during larval development and their morphology is unchanged in the mutants, we conclude that Ral plays a role specifically in wrapping glia in the regulation of membrane growth. Finally, we detected no structural defects in the lamella of *ral* mutants (Figure 2E), akin to perineurial (which secrete some of the ECM components of the lamella), and to subperineurial glia. Similar results were obtained for the nerves that exit the VNC (Supplementary Figure 2). In summary, our structural analysis reveals that Ral GTPase plays a role specifically in the regulation of wrapping glia morphology.

During larval development, wrapping glia do not divide but their membranes need to grow to accompany the ∼100 times increase in larval size (Hilchen et al. 2013). Given that this is a continuous developmental process, the defects observed in *ral* mutants (Fig. 2D) can reflect a failure to grow during development or alternatively reflect a collapse of pre-formed membranes. To distinguish between these possibilities, we analyzed wrapping glia morphology from late 1^st^-instar, through late 3^rd^-instar larvae (Figure 3). Our data clearly shows that the wrapping glia membrane does not grow during development. Contrary to the controls where, by late 1^st^-instar, a uniform membrane that covers the length of the nerve is visible (Figure 3A), in the mutants the wrapping glia membrane remains as a thin thread throughout larval development, without signs of ever growing or ensheathing the axons (Figure 3B-C). Therefore, we conclude that Ral GTPase is necessary for the growth of wrapping glia membrane during development, a process critical for axonal ensheathing and efficient synaptic conduction.

**Figure 3.**
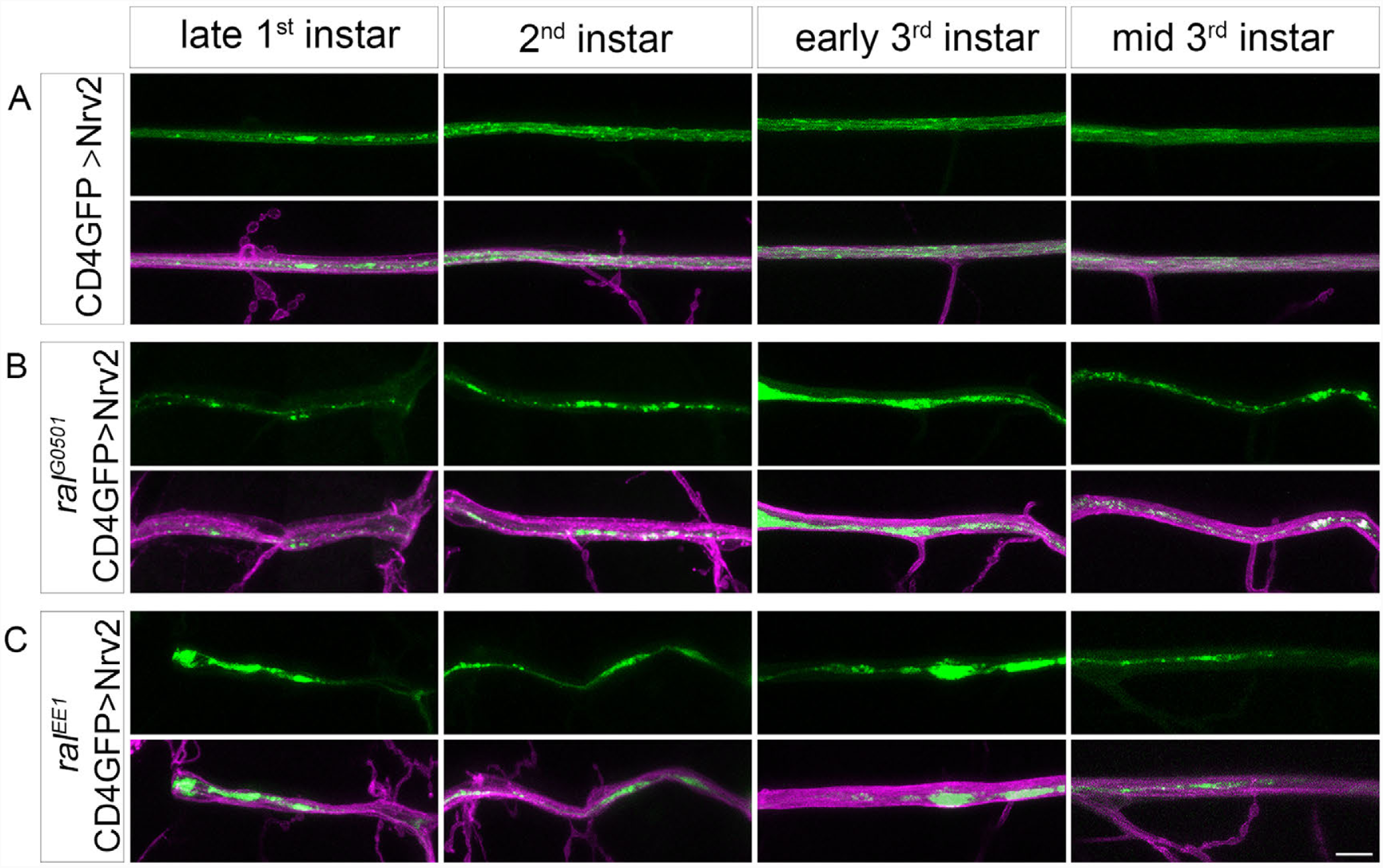
Ral GTPase is required for wrapping glia development, not maintenance. **(A-C)** Confocal images of wrapping glia membranes labeled with CD4-GFP (green, Nrv2-Gal4>UAS-CD4-GFP) and the nerve labelled with HRP (magenta). Images in **(A)** *w*^*1118*^, **(B)** *ral*^*G0501*^ and **(C)** *ral*^*EE1*^ correspond to wrapping glia morphology in different larval stages. The defects in wrapping glia (in green) in *ral* mutants are visible from 1^st^ instar larval stage and are persistent throughout larval development. Scale bar: 10µm.

### ral mutants have abnormal ultrastructural nerve profiles and defective septate junctions

While it is clear that Ral is necessary for wrapping glia development, it is paradoxical that defects in a glia subtype results in widespread thickening of ISNs. The increase in nerve thickness can result from defective axonal fasciculation or from BNB problems (Leiserson et al. 2000; Yu et al. 2000; Baumgartner et al. 1996; Luong et al. 2018). To examine the ultrastructural features of nerve bundles with respect to the presence, distribution and organization of its cell types, we used transmission electron microscopy (TEM). We dissected 3^rd^-instar larvae keeping the brain and nerve bundles intact, and obtained ultrathin sections of the segmental nerves between 2-10 μm after the exit of the VNC, as described (Matzat et al. 2015). TEM images of WT and *ral* mutant larvae revealed that the inner structure of the bundle in the mutants had a high incidence of large electron-transparent regions (ETR), occupying an average of 6.55±1.49% in *ral*^*G0501*^ and 10.28±2.31% in *ral*^*EE1*^ of the nerve cross-area comparing with 0.93±0.33% in controls (Figure 4A-C, and Supplementary Figure 3). In addition to the increased incidence of ETRs, the ultrastructural analysis of the nerves supports the notion that wrapping glia are underdeveloped in *ral* mutants (cyan). When possible, we counted the number of axonal profiles present in controls and in mutants and observed that the numbers were identical between genotypes, and correspond to what has been reported (Leiserson et al. 2000; Matzat et al. 2015). This data is also consistent the immunofluorescence results where we observe normal neuronal patterning. However, the observation of these ETRs raised the possibility that the BNB may be compromised and that this phenotype represents a breach of this barrier.

**Figure 4.**
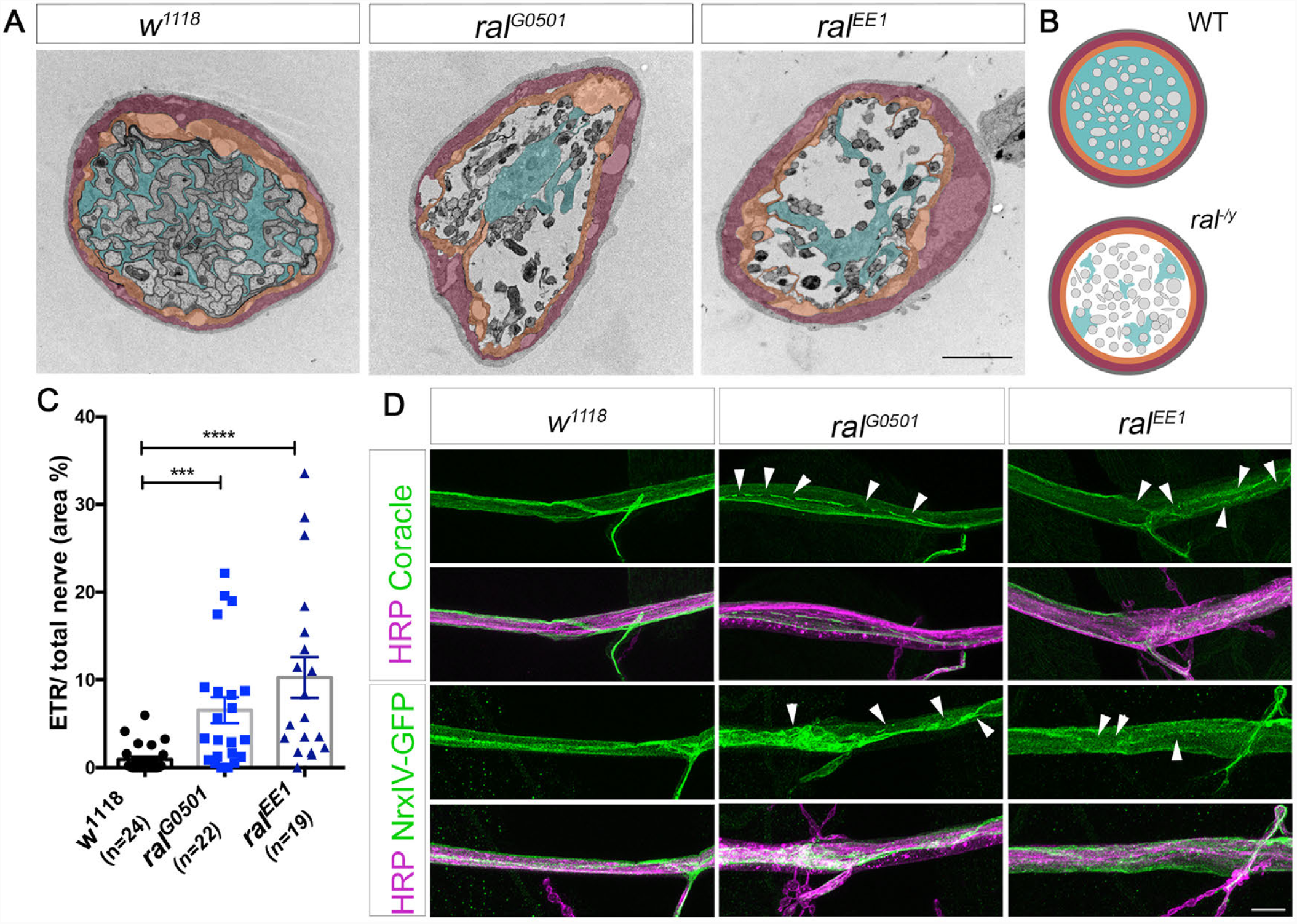
*ral* mutants have abnormal peripheral nerve ultrastructure and disrupted septate junction markers. **(A)** Representative examples of transmission electron microscopy (TEM) images of transversal cut of segmental nerves within 2-10μm from the exit of the VNC of 3^rd^-instar larvae. ETR within the wrapping glia layer is larger in the nerves of *ral*^*G0501*^ and *ral*^*EE1*^. These areas can result from the absence of wrapping glia or are probably filled with haemolymph. Scale bar: 2μm **(B)** Schematic representation of segmental nerves displaying the defects observed with TEM. Above is represented a WT nerve and below a mutant nerve. Light grey – axon, light blue – wrapping glia; orange –subperineurial glia; magenta – perineurial glia and dark grey – neural lamella. **(C)** Quantifications of the area of the electron-transparent regions (ETR)/total nerve area. N= number of nerves analysed. Non-parametric ANOVA, Kruskal Wallis with Dunn’s multiple comparison test was used. ****p<0.0001 ***p<0.001. **(D)** Representative examples of the ISN labelled with Coracle (green) or Neurexin IV-GFP (NrxIV-GFP, green), components of SJs, and with HRP (magenta) to label the ISN membrane in *w*^*1118*^, *ral*^*G0501*^ and *ral*^*EE1*^ in 3^rd^-instar larvae. Arrows indicate examples of disruptions in distribution of Coracle or NrxIV-GFP. Scale bar: 10μm

The BNB is the cell barrier responsible for keeping the larval haemolymph from entering the nerve, and SJ are the cellular structure responsible for the formation of the BNB (Stork et al. 2008). SJs are homologue structures to vertebrate tight junctions (TJ) and are composed of several proteins, including Neurexin-IV (NrxIV), and Coracle (Banerjee et al. 2006). Interestingly, TJ formation has been shown to depend on Ral function (Hazelett et al. 2011), and it is therefore plausible that *Drosophila* Ral GTPase also plays a role in PNS SJ establishment. Given that SJ mutants die in late-stage embryos (Hall & Ward 2016; Baumgartner et al. 1996), it is unlikely that SJs fail to form completely in *ral* mutants, but it is possible that milder defects are present. To address SJ integrity we recurred to antibodies raised against Coracle and to a GFP-tagged protein trap in the Nrx-IV *locus*, which allowed us to visualize SJs in controls and in *ral* mutants. As it can be seen in Figure 4D, in controls both Coracle and NrxIV-GFP appear as a continuous line that represents the auto-SJ formed by subperineurial glia. In *ral* mutants, even though Coracle and Nrx-IV do localize to the membrane, their distribution is uneven, exhibiting several bifurcations and interruptions, not visible in controls (Figure 4D, arrowheads). These anomalies can represent places where SJs are abnormally assembled and where the BNB may be breached and leaky. In conclusion, both TEM and immunofluorescence suggest that in addition to defects in wrapping glia growth, the BNB may be compromised in *ral* mutants.

### ral mutants have locomotor defects

Changes in axonal wrapping have been associated with increased spike propagation times (Ghosh et al. 2013). As wrapping glia are responsible for insulating axons, defects in wrapping glia membrane growth should lead to reduced axonal insulation, altered electrical conduction and, ultimately, to locomotor problems. To assess whether this was the case in *ral* mutants, we performed a larval locomotion assay in which wandering 3^rd^-instar larvae were filmed for 5 minutes while freely wandering in an agar arena, and their positions were tracked with IdTracker Software (rez-Escudero et al. 2014). WT larvae displayed the expected bimodal crawling behaviour, alternating between active crawling and reorientation events, and crawled in average 134,6±7,7mm in 5 minutes, with an average speed of 0,52±0,03mm/s. *ral* mutants also displayed the bimodal pattern, but crawled less and at slower speed: during the 5 minutes analysed *ral*^*G0501*^ and *ral*^*EE1*^ larvae crawled in average 92,5±5,7mm and 88.2±4.4mm, at an average speed of 0,37±0,02mm/s and 0,39±0,02mm/s, respectively (Figure 5A-C). Interestingly, we observed that, at times, *ral* mutant larvae seemed to drag their posterior body-half, as the peristaltic contractions appeared incomplete, losing power along the anterior-posterior axis (compare WT and mutant larvae in movies 1-3). To uncover if the differences between WT and the mutants were due to the mutants crawling slower in the active crawling phase or spending more time in reorientation events, we analysed the speed of long uninterrupted forward crawls. WT larvae had an average active crawling speed of 0,67±0,07mm/s, while *ral*^*G0501*^ and *ral*^*EE1*^ mutants performed fewer long forward runs and crawled at significantly lower speeds: 0,38±0,02mm/s and 0,38±0,04mm/s, respectively (Figure 5D). This data is consistent with our hypothesis of signal dissipation in axons, suggesting that motor axons in peripheral nerves lacking fully developed wrapping glia fail to efficiently propagate action potentials, leading to locomotor deficits. However, given that this assay is performed in *ral* mutant larvae, where the gene is mutated in all tissues, we cannot conclude if this defective behaviour derives from Ral being required in glia or elsewhere.

**Figure 5.**
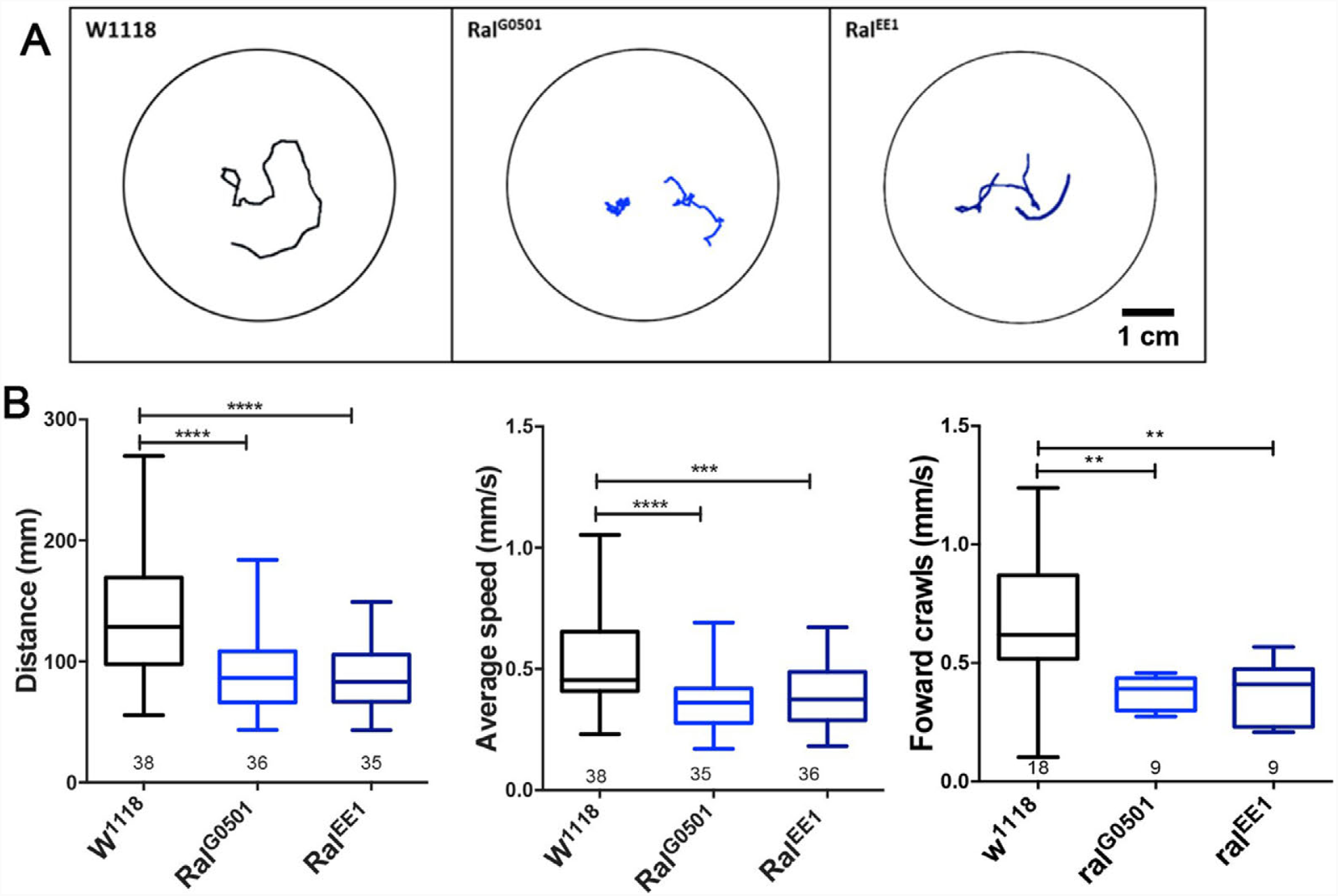
*ral* mutants have impaired locomotion. **(A)** Representative examples of larvae trajectories after 5min of crawling in an open field arena for the genotypes *w*^*1118*^, *ral*^*G0501*^ and *ral*^*EE1*^. **(B)** Quantification of the total distance travelled by 3^rd^-instar larvae of the indicated genotypes. *ral* mutants have impaired locomotion. Scale bar: 1 cm. **(C)** Quantification of the average velocity during the entire 5min session of the indicated genotypes. **(D)** Quantification of the speed of forward crawls. N represents the number of animals analysed. Ordinary one-way ANOVA with Bonferroni multiple comparison test was performed. ****p<0.0001 ***p<0.001.

### Ral GTPase is required in wrapping glia to promote growth

We identified Ral GTPase as necessary for wrapping glia development, but whether it is required cell autonomously cannot be concluded from our mutant analysis. To test this, we used a knock-down strategy using RNAi to reduce the levels of Ral GTPase in a cell-type specific manner, using the same Gal4 lines as before, followed by ISN thickness analysis as readout for developmental alterations. As controls, we used the UAS-Ral-IR, and each of the Gal4 lines, crossed with WT. Ral RNAi (Ral-IR) in all glia, using Repo-Gal4, leads to widespread thickening of the ISN with clear signs of defasciculated axons (Figure 6A-C). Likewise, using the wrapping glia specific driver (Nrv2-Gal4) to reduce Ral, also induces an ISN significantly thicker than the controls (Figure 6A-C), implicating that Ral in wrapping glia is required for the regulation of nerve bundle morphology. Interestingly, even though both all glia and wrapping glia Ral-IR results in ISN thickening, we also observe the appearance of swellings along the ISN, but only in all glia knock down (Supplementary Figure 4). This result corroborates the notion that Ral, in addition to regulating wrapping glia development, also plays additional roles in other subtypes, including in subperineurial glia during BNB formation.

**Figure 6.**
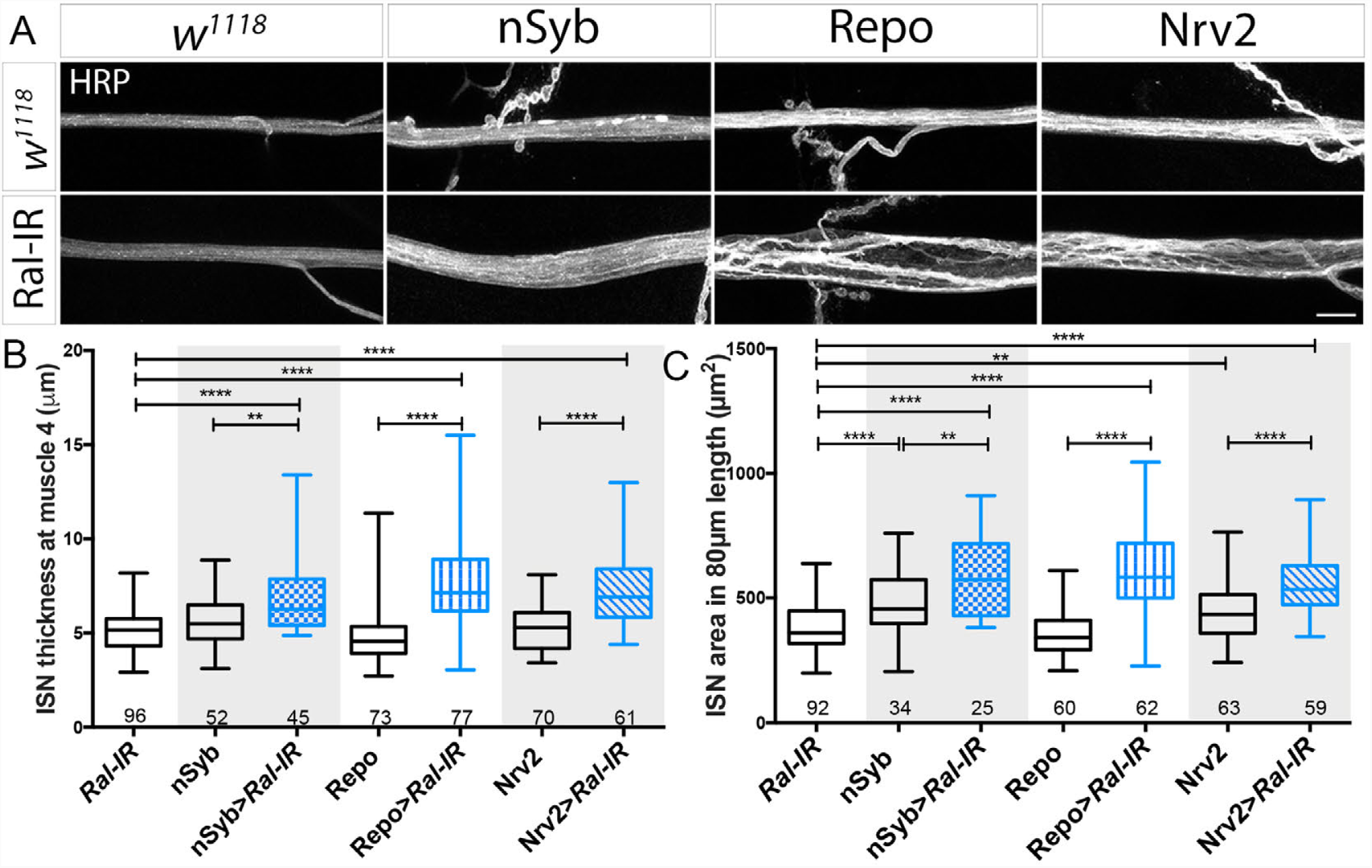
Ral GTPase reduction in neurons and glia lead to nerve thickenings. **(A)** Representative examples of ISN at m4 exit after cell-type specific Ral-IR using each of the Gal4 drivers indicated. Ral knockdown was achieved by crossing UAS-Ral-IR with tissue specific drivers (nSyb-Gal4>neurons, repo-Gal4> pan-glia and Nrv2-Gal4>wrapping glia). As control, *w*^*1118*^ flies were crossed with UAS-Ral-IR and with each of the Gal4 drivers. ISN thickness was used as readout. **(B)** Quantifications of the width of the ISN at m4 exit point of the indicated genotypes. Non-parametric ANOVA, Kruskal-Wallis with Dunn’s multiple comparisons test was used, ****p<0.0001, **p<0.005. **(C)** Quantification of the nerve area, per 80μm of ISN in the same genotypes as in (B). Ordinary one-way Anova test was used. ****p<0.0001, **p<0.005. All comparisons were performed between the Ral-IR, Gal4 control and experimental group. Numbers on x-axis represent total number of ISNs quantified from segments A2-A4. Scale bar: 10μm.

To our surprise, we find that ISN thickness is also increased when Ral is reduced in neurons using the panneuronal driver nSyb-Gal4 (Figure 6A-C), suggesting that Ral is also required in neurons to mediate some aspect of ISN development. Given that mutants also have some degree of defasciculation (Figure 2A and Supplementary Figure 1), we hypothesized that *ral* could be important in the regulation of adhesion. Of particular interest is Fasciclin-II (FasII) because it is one of the most abundant adhesion proteins present in motor neurons and that promotes neuron-neuron and neuron-glia adhesion (Wright & Copenhaver 2001). However, quantification of FasII levels by immunofluorescence showed no differences between wild-type and *ral* mutants (Supplementary Figure 5A-B). Despite this, it is still possible that adhesion molecules other than FasII are altered in *ral* mutants and are responsible for the observed axonal defasciculation. With this idea in mind, we asked if the defasciculation phenotype could be rescued by promoting adhesion. To test this, we expressed full length FasII in motor neurons, using the motor neuron-specific driver OK6-Gal4 and calculated ISN thickness (Supplementary Figure 5C-D). We found that this manipulation was able to completely rescue the thick nerve phenotype present in the mutants, supporting our hypothesis that adhesion may contribute to the defects, but are not due to alterations in FasII expression levels. This data revealed that the thickening of the ISN results from a combination of factors including lack of ensheathment, adhesion, SJ integrity, and fasciculation.

To further validate the cell type specific findings from the RNAi experiments, we performed cell-type specific rescues in the *ral* mutants. For this, we expressed UAS-HA-tagged Ral (RalHA) under the control of cell-type specific Gal4 lines, in the background of each of the *ral* mutants. Quantification of nerve thickness was used as readout for rescue. Neuronal rescues of Ral give rise to a nerve significantly thinner than *ral* mutants (Figure 7A,C). This result is consistent with the RNAi data (Figure 6), and may indicate that reintroduction of Ral in neurons rescues a still unidentified adhesion factor. When Ral is expressed in all glia using Repo-Gal4, nerve thickness is rescued (Figure 7A, C). We also quantified the nerve area in 80μm length of ISN and observed that putting back Ral in all glia rescues the phenotype in both *ral* mutants (Figure 7D). The slight discrepancies in rescue capacity of the two mutants probably reflect some intrinsic differences between the lines or in the genetic backgrounds. Reintroduction of Ral exclusively in the wrapping glia of *ral* mutant animals using Nrv2-GAL4, also rescues the enlarged nerve phenotype but not always to wild-type levels (Figure 7A, C). Importantly, because the rescue construct has an HA-tag we can assess where Ral is being expressed. Because Ral is known to localize to the plasma membrane irrespective of its nucleotide binding state (Teodoro et al. 2013), we can extrapolate the morphology of wrapping glia. Our results clearly show that Ral expression in wrapping glia leads to a complete rescue of wrapping glia morphology (Figure 7B). Even in cases where there is still a thick nerve (HRP vs Ral-HA staining in Figure 7B), wrapping glia growth was rescued. This result indicates that Ral is necessary in wrapping glia to regulate its own growth. Additionally, expression of Ral-HA in only wrapping glia does not rescue the SJ marker Coracle distribution (data not shown), suggesting that Ral cannot regulate SJ formation non-cell autonomously. In fact, the observation in this experiment that there are enlarged ISN, despite having normal wrapping glia, indicates that the problems with the BNB are independent of the wrapping glia. In summary, a balance of factors in neurons and in glia must be tightly coordinated to regulate ISN development and function.

**Figure 7.**
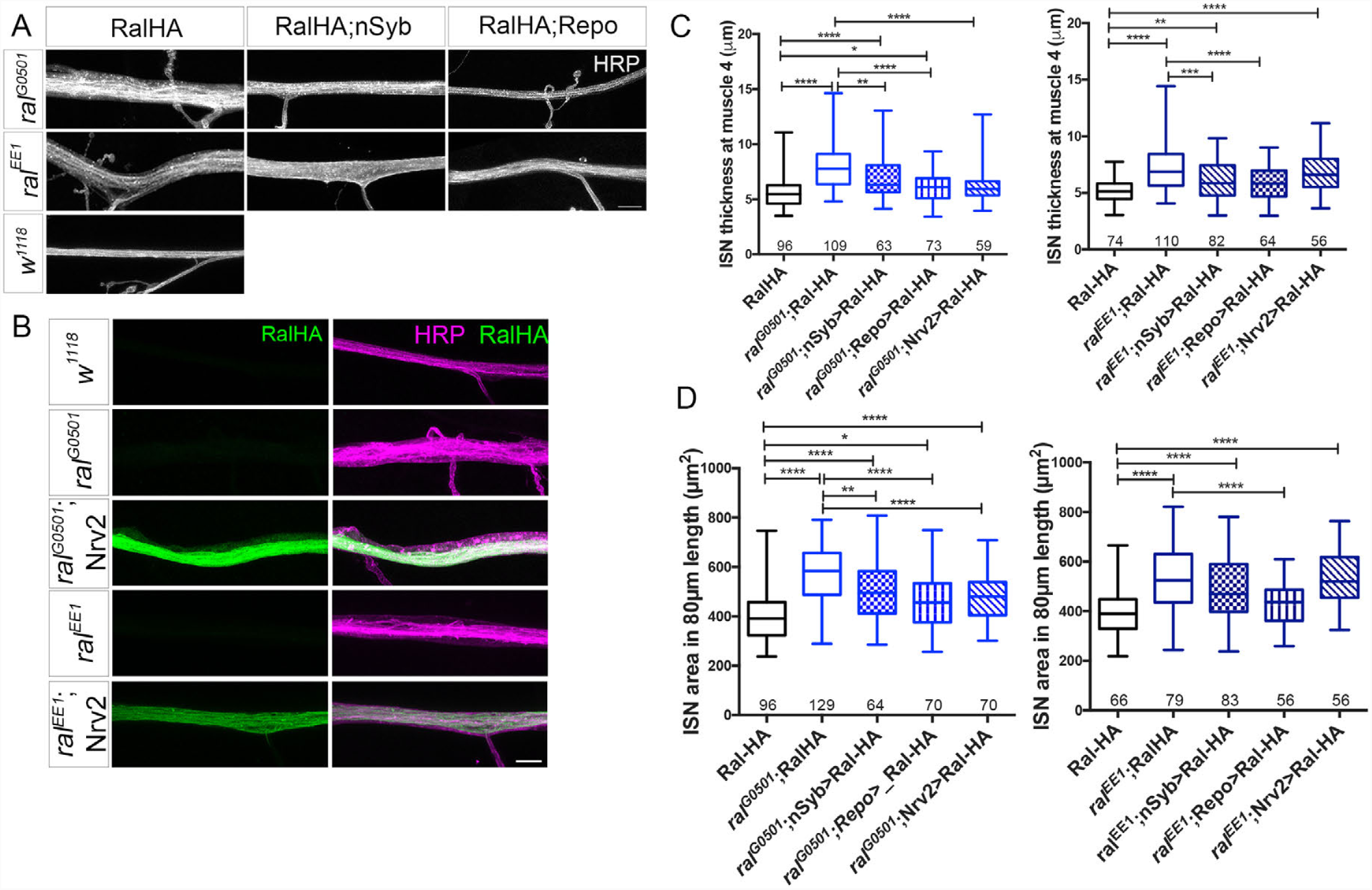
Ral GTPase is required cell autonomously to regulate wrapping glia development and growth. Representative examples of rescue experiments of *ral* mutants expressing UAS-RalHA under the control of tissue specific drivers (nSyb-Gal4>neurons and repo-Gal4> pan-glia), in *ral*^*G0501*^ and *ral*^*EE1*^ background. HRP (grey) labeling was used to visualize the ISN. Full genotypes from left to right, top to bottom, are: UAS-RalHA | *ral*^*G0501*/Y^; UAS-RalHA | *ral*^*G0501*/Y^; UAS-RalHA>nSyb-Gal4 | *ral*^*G0501*/Y^; UAS-RalHA>Repo-Gal4 | *ral*^*EE1/*Y^; UAS-RalHA | *ral*^*EE1/*Y^; UAS-RalHA>nSyb-Gal4 | *ral*^*EE1/*Y^; UAS-RalHA>Repo-Gal4 | *w*^*1118*^Representative examples of rescue experiments of *ral* mutants expressing UAS-RalHA under the control of a wrapping glia specific driver (Nrv2-Gal4>WG) in *ral*^*G0501*^ and *ral*^*EE1*^ background. Labeling of Ral-HA (green) using anti-HA antibody allows visualization of WG morphology. ISN membrane labeled with HRP (magenta). Full genotypes, top to bottom, are: UAS-RalHA | *ral*^*G0501*/Y^; UAS-RalHA | *ral*^*G0501*/Y^; UAS-RalHA>Nrv2-Gal4 | *ral*^*EE1*/Y^; UAS-RalHA | *ral*^*EE1*/Y^; UAS-RalHA>Nrv2Gal4. Scale bar: 10μm. **(C)** Quantification of ISN thickness, at m4 exit point. **(D)** Quantification of the nerve area, per 80μm of nerve. Non-parametric ANOVA, Kruskal Wallis with Dunn’s multiple comparisons test were used. ****p<0.0001, ***p<0.0005, **p<0.005, *p<0.05. Numbers on x-axis represent the total number of ISNs quantified from segments A2-A4.

### Ral GTPase regulates nerve bundle development via the Exocyst complex

Several different pathways have been reported to act downstream of Ral GTPase, including the exocyst complex, RalBP1 and Filamin (Moghadam et al. 2017; Shirakawa & Horiuchi 2015). Ral interaction with RalBP1 is mostly involved in endocytosis, Filamin and Ral act as scaffold/synaptic organizers (Lee & Schwarz 2016), and the Ral/Exocyst pathway has been shown to be required for membrane addition of postsynaptic membranes and for the polarized trafficking of vesicles to exocytic places (Teodoro et al. 2013; Armenti et al. 2014; Balasubramanian et al. 2010; Wang et al. 2004; Hase et al. 2009). From these interactors, the exocyst is an excellent candidate to mediate Ral-dependent wrapping glia growth. If the exocyst complex regulates nerve bundle morphology, mutants should have a phenotype similar to the one observed in *ral* mutants. In *Drosophila*, most exocyst mutants die as 1^st^-instar larvae but a mutation induced by the insertion of a P-element in *sec8 locus* (*sec8*^*P1*^) results in a hypomorphic mutants that survive to 3^rd^-instar stages (Liebl et al. 2005). We analyzed the thickness of the ISN in this mutant and in a control line where the P-element was precisely excised (*sec8*^*revs*^). Our data shows that control larvae (*sec8*^*revs*^) have an ISN that is 5.67±0.13μm, which is significantly different from *sec8*^*P1*^where the ISN is thicker being 7.08±0.13μm (Figure 8A-B). Therefore, like *ral* mutants, *sec8*^*P1*^ larvae have thicker ISNs than its control line.

**Figure 8.**
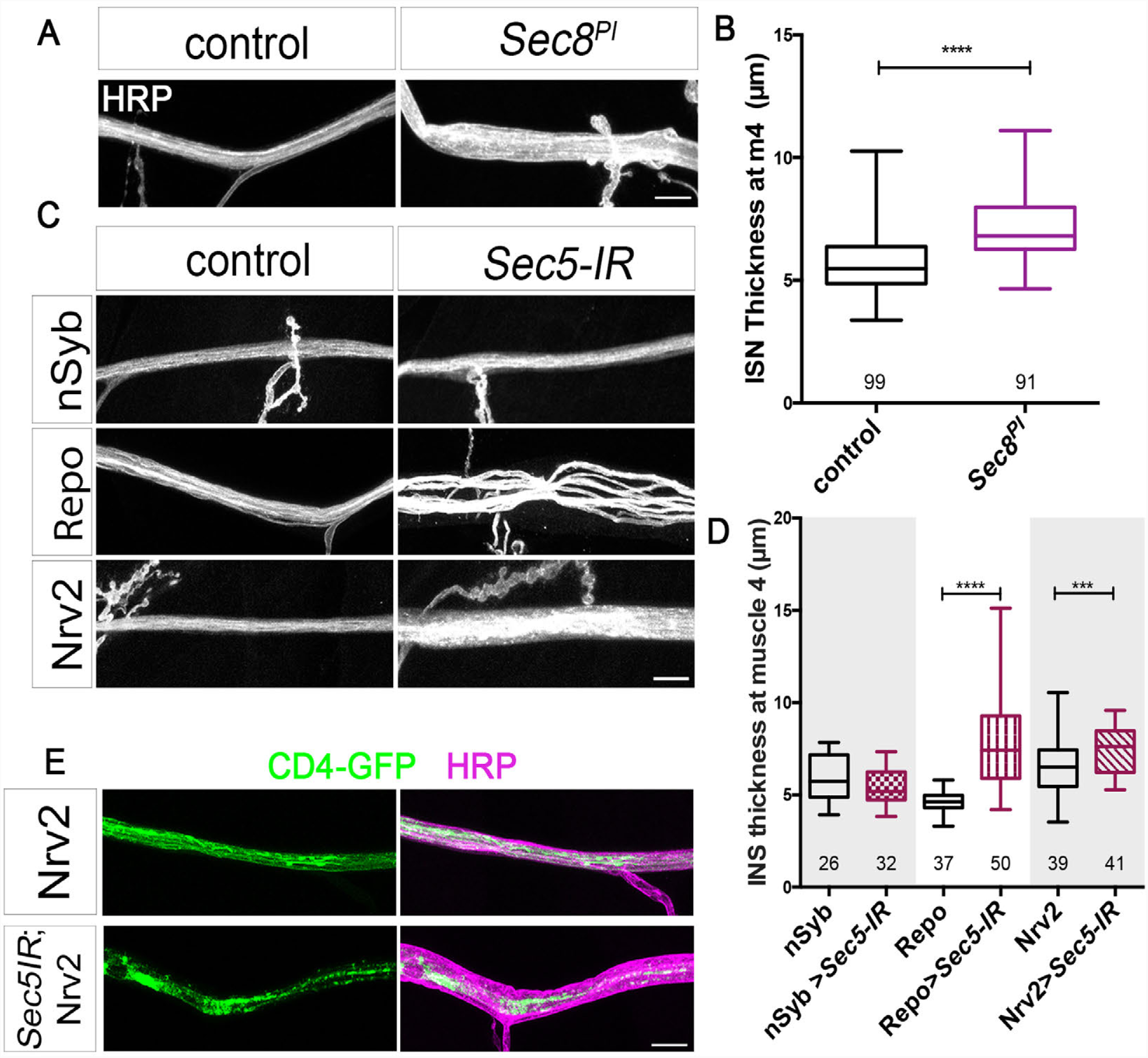
Reducing the levels of the exocyst – a Ral GTPase effector, mimics *ral* mutant phenotypes. **(A)** Representative examples of the ISN labeled with HRP (grey) to outline the nerve in control (*sec8*^*revs*^) and in *sec8* hypomorphic mutants (*sec8*^*P1*^*/sec8*^*P1*^). Scale bar: 10μm. Quantifications of the width of the ISN at m4 exit. Unpaired Mann-Whitney test was used ****p<0.0001. **(C)** Cell-type specific knockdown of Sec5 (Sec5-IR) using RNAi was achieved by crossing UAS-Sec5-IR with tissue specific drivers (nSyb-Gal4>neurons, Repo-Gal4>pan-glia and Nrv2-Gal4>wrapping glia). As control, *w*^*1118*^ flies were crossed with each of the Gal4 drivers. Representative examples of ISN at muscle 4 exit after Sec5-IR expression with each of the Gal4 drivers indicated. ISN thickness was used as readout. **(D)** Quantifications of the width of the ISN at m4 exit point of the indicated genotypes in μm. Scale bar: 10μm. Mann-Whitney test was used between Gal4 control and experimental group. ****p<0.0001***p<0.001. **(E)** Morphological analysis of wrapping glia in Nrv2>Sec5-IR was performed by co-expressing CD4-GFP together with the RNAi. Representative examples of the ISN labeled with CD4-GFP (green) and HRP (magenta) to outline the nerve in control and Nrv2>Sec5-IR. Scale bar: 10μm. Numbers on x-axis represent the total number of ISNs quantified from segments A2-A4.

To address which cell type is responsible for this defect, we used cell-type specific Gal4 lines to do RNAi against another exocyst subunit - Sec5, which is known to bind directly to Ral and reported to promote membrane growth in other systems (Teodoro et al. 2013). When RNAi against Sec5 (Sec5-IR) was expressed in neurons, using nSyb-Gal4, there was no effect on the morphology of the bundle (Figure 8C-D, nSyb=5.92±0.25μm, nSyb>Sec5-IR=5.41±0.17μm). However, expression of Sec5-IR in all glia resulted in a very thick nerve, with extreme signs of axonal defasciculation, highly reminiscent of *ral* mutants (Figure 8C-D, Repo=4.64±0.10μm, Repo>Sec5-IR=8.02±0.38μm). When Sec5-IR was expressed exclusively in wrapping glia, using Nrv2>Sec5-IR, the nerve was also significantly thicker than the Gal4 control (Figure 8C-D, Nrv2=6.40±0.25μm, Nrv2>Sec5-IR=7.48±0.19μm), resembling Ral-IR in wrapping glia (Figure 6A). Importantly, labeling the membrane of wrapping glia (with CD4-GFP) in Sec5-IR shows that reduction of Sec5 in wrapping glia impairs their growth, akin to *ral* mutants (Figure 8E, compare to Figure 2D). Rather than completely ensheathing the axons, wrapping glia in Nrv2>Sec5-IR did not fill the entire nerve area and were thinner and less regular than controls.

If Ral and the exocyst complex are in the same genetic pathway, heterozygous mutations of these genes should genetically interact and give rise to a thick nerve phenotype. We postulated that mutating a single exocyst component would probably be insufficient to reveal an interaction between this 8-protein complex and Ral GTPase. Therefore, we tested if heterozygous mutations in *ral* genetically interacted with double-heterozygous mutations in two exocyst components - Sec5 and Sec6. For this experiment, we used a line where null mutations in the *loci* of *sec5* (*sec5*^*E10*^) and in *sec6* (*sec6*^*Ex15*^) were recombined and tested in heterozygosity. This double mutant was tested in combination with heterozygous *ral* mutations, *ral*^*G0501*^ and *ral*^*EE1*^. To assess genetic interactions, we measured ISN thickness in the different heterozygous combinations. Strikingly, in contrast with the single and double mutants, the triple heterozygous of *ral, sec5*, and *sec6* exhibits a significantly thicker nerve (compare Figure 1C and Figure 9). Heterozygous *ral*^*G0501*^ and *ral*^*EE1*^ were indistinguishable from wild-type (*w*^*1118*^). Double heterozygous mutations of *sec5,sec6* have a small but significant increase in nerve thickness compared to wild-type, but that is strongly potentiated by the introduction of one mutated allele of the *ral* gene (Figure 9B). In conclusion, this data demonstrates that Ral GTPase regulates nerve bundle development via the exocyst complex.

**Figure 9.**
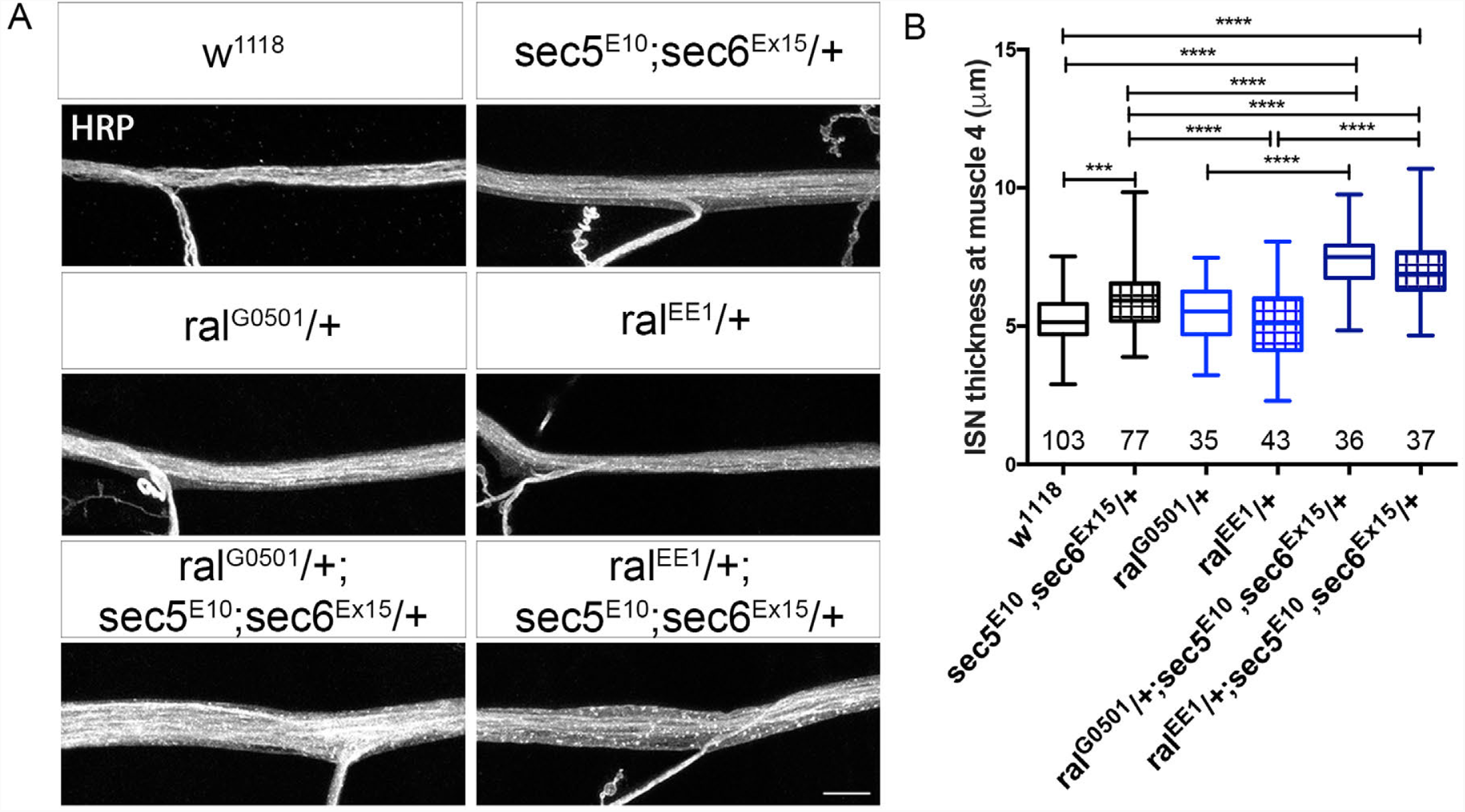
Genetic interactions show that Ral GTPase acts via the exocyst complex in the regulation of peripheral nerve thickness. **(A)** Representative examples of the ISN labeled with HRP (grey) to outline the nerve in the following genotypes represented left to right, top to bottom: control (*w*^*1118*^) | *sec5*^*E10*^, *sec6*^*Ex15*^*/+* | *ral*^*G0501*^/+ | *ral*^*EE1*^/+ | *ral*^*G0501*^/+; *sec5*^*E10*^, *sec6*^*Ex15*^*/+* | *ral*^*EE1*^/+; *sec5*^*E10*^, *sec6*^*Ex15*^*/+*. Scale bar: 10μm. **(B)** Quantification of the width of the ISN at m4 exit of the indicated genotypes in μm. *ral* and exocyst heterozygous mutations interact genetically giving rise to a thicker ISN. Numbers on x-axis represent total number of ISNs quantified from segments A2-A4. Parametric one-way ANOVA, with Bonferroni multiple comparisons test was used. ***p<0.0005, ****p<0.0001.

## Discussion

Axonal ensheathment is a fundamental biological process that allows fast impulse conduction and the correct functioning of the nervous system (Saab & Nave 2017; Seidl 2014). Despite its importance, the molecular players responsible for the regulation of this process are ill defined. Our study identifies Ral GTPase and the exocyst complex as novel factors required for wrapping glia growth and development *in vivo* (Figure 10). Through mutant analysis, cell-type specific RNAi and rescue experiments, we discovered that Ral GTPase plays a specific role in the regulation of wrapping glia development, and that it is both sufficient and necessary for the growth of this glial subtype. We also discovered that *ral* mutants have locomotion defects and show signs of a disrupted BNB and axonal defasciculation. The defects observed in exocyst mutants and RNAi suggest that, at the ISN, glia-dependent functions are mediated via the exocyst, while neuronal fasciculation functions are not. Based on our findings, we propose a mechanism whereby activation of Ral GTPase in wrapping glia induces the recruitment of the exocyst, probably via Sec5 interaction, targeting vesicles to the membrane. These vesicles can directly contribute to membrane growth, but can also carry a wrapping glia -specific secreted factor, that can signal wrapping glia growth (Figure 10). Our study identifies a new *in vivo* mechanism by which glia regulate axonal ensheathing, providing novel insights onto the molecular basis of nerve development.

**Figure 10.**
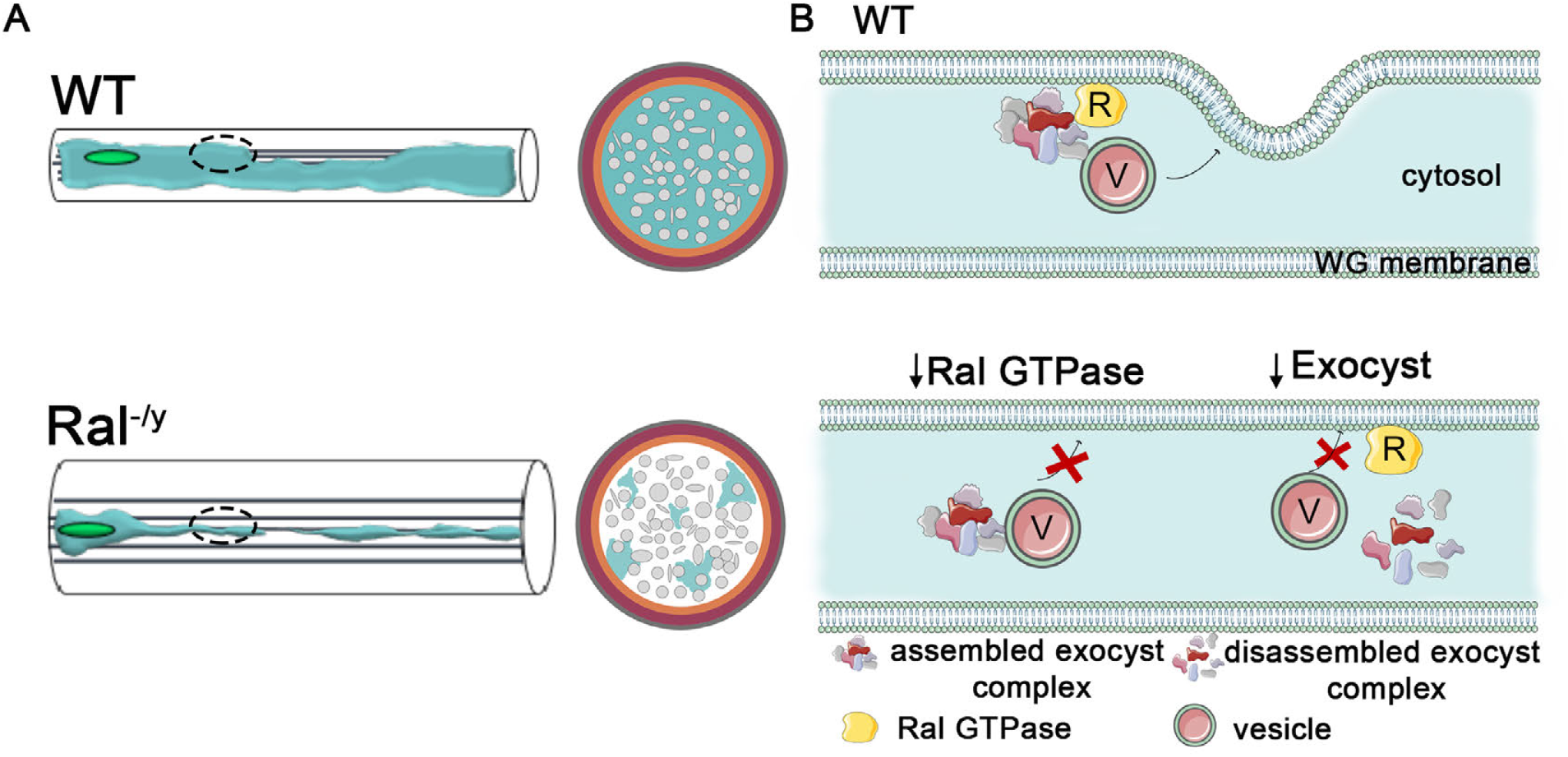
Model of how Ral GTPase and the exocyst complex regulate wrapping glia development. **(A)** Schematic of a wild-type (WT-top) and of a *ral* mutant (*ral*^*-/Y*^-bottom) ISN represented longitudinally (left) and in a cross section (right). In *ral* mutants wrapping glia are underdeveloped. Colors represent the following structures: light grey–axons, light blue– wrapping glia; orange–subperineurial glia; magenta–perineurial glia and dark grey– neural lamella **(B)** Cartoon of suggested model for Ral GTPase and exocyst-dependent participation in wrapping glia development. We propose that in wild-type <u>(top)</u>, Ral GTPase at the plasma membrane leads to the engagement of the exocyst complex and consequent targeting of vesicles to the membrane. Fusion of these vesicles can promote wrapping glia growth directly or can carry a wrapping glia-derived growth signal that will then regulate their growth. In the absence of Ral <u>(bottom left)</u>, exocyst recruitment is compromised and vesicles cannot fuse. For simplicity, we represent the exocyst complex assembled in the absence of Ral, but our data does not address assembly status directly. In the <u>bottom right</u>, we schematize the situation where subunits of the exocyst complex are reduced. Here, even though Ral GTPase can localize to the membrane, the exocyst complex cannot assemble correctly, nor can secretory vesicles be recruited to the membrane, resulting in defective membrane addition and underdeveloped wrapping glia.

We identified Ral and exocyst as factors required for wrapping glia membrane growth. This pathway has been implicated in membrane addition in other cellular contexts, including postsynaptic growth (Teodoro et al. 2013; Patrício-Rodrigues & Teodoro 2018), nanotube formation (Hase et al. 2009) and in growth phases during cell division (Murthy et al. 2010; Holly et al. 2015; Giansanti et al. 2015). While the Ral/Exocyst pathway is clearly a mechanism widely used for growth regulation, our results indicate that it is not simply a default growth signal, since subperineurial glia size was unchanged in the mutants and this glia subtype also depends on hypertrophic growth during larval development. Therefore, one (or more) yet-unknown factor(s) must confer this specificity.

Previous studies have identified molecules required for wrapping glia development, including Vein (Matzat et al. 2015), the homologue of Neuregulin-1, and Akt (Lavery et al. 2007) - two genes that also play a role in mammalian myelination (Michailov 2004; Domenech-Estevez et al. 2016). Additionally, mutations in *fray* (a serine/threonine kinase) (Leiserson et al. 2000), *khc1* (Schmidt et al. 2012), *lace* (a serine palmitoyltransferase) (Ghosh et al. 2013), integrin signaling (Xie & Auld 2011) and Laminin (Petley-Ragan et al. 2016) also result in defective wrapping glia development. Some of these pathways can potentially intersect with Ral/Exocyst function. Namely, the Neuregulin-1 homologue Vein has been shown to be required in wrapping glia to mediate its own development, and non-cell autonomously to induce SJ formation in subperineurial glia. Knowing that the exocyst is required for the correct trafficking and secretion of a different EGFR ligand, Gurken (Murthy & Schwarz 2004), and Ral has been implicated in EGFR activity (Lu et al. 2000), it is conceivable that the Ral/Exocyst pathway can be required for the secretion of Vein, which would then instruct wrapping glia growth and development. Nonethless, contrary to Vein, Ral does not regulate SJs non-cell autonomously, since reintroducing Ral in wrapping glia does not rescue the distribution of Coracle, one of the subperineurial glia SJ components (data not shown). However, given that we observed disrupted SJs in *ral* mutants, it is probable that Ral functions directly in subperineurial glia to regulate SJ assembly. In fact, RalA via the exocyst, has been shown to be required for normal tight junction assembly and barrier function in MDCK epithelial cells (Hazelett et al. 2011), making this pathway a candidate for regulating BNB formation in the *Drosophila* PNS.

In addition to interacting with Ral GTPase, the exocyst is also an effector to several Rab GTPases (Wu & Guo 2015). The wrapping glia morphology defects observed in *khc* mutants were correlated with trafficking problems of Rab21 and Rab30 and it is therefore possible that the exocyst can interact with these Rab GTPases (Schmidt et al. 2012). Interestingly Rab21 has been implicated in integrin signaling (Pellinen et al. 2008) and integrins are also important for wrapping glia development, where βPS integrin reduction results in underdeveloped wrapping glia (Xie & Auld 2011). In vertebrates, RalA mediates integrin-dependent membrane raft exocytosis and growth signaling (Balasubramanian et al. 2010). Thus, it is plausible that Ral/exocyst and integrins cooperate during axonal ensheathment, possibly by being involved in the regulation of wrapping glia growth and maintenance. However, we observed normal βPS localization in *ral* mutants (data not shown), making it less likely that integrin signaling is the origin of the wrapping glia defects.

Even though invertebrates do not produce myelin, it is possible that the genes important for accomplishing such an important function as axonal ensheathing, are conserved, even if with some layers of adaptation. The fact that in vertebrates Ral and exocyst are present in myelin (Anitei 2006; Anitei et al. 2009) and that Sec8 has been implicated in secretory events that promote myelin sheath growth (Bolis et al. 2009), support this idea. In this work, we found that cell autonomous manipulation of the levels of Ral GTPase in wrapping glia leads to severe defects in axon ensheathment and *ral* mutants have strong locomotor deficits, supporting an important physiological role of Ral GTPase for the control of movement. Defects in axonal wrapping are the main underlying cause of the symptoms present in patients with multiple sclerosis (Pan & Chan 2017; Saab & Nave 2017). While myelination in the PNS has long been appreciated as required for movement, more recent studies have shown that in neurological disorders of the CNS, like Alzheimer’s Disease (AD), there are signs of myelin loss in the white matter, even in pre-clinical symptoms (Saab & Nave 2017), making the understanding of the mechanism by which ensheathing is regulated ever more relevant. Currently, no therapies are available that can induce any adjustment to the levels of axonal wrapping, which precludes the treatment of disorders that have either too little or too much myelin. Identification of the specific molecules that regulate axonal wrapping in the PNS and CNS can help finding new genes whose function can be used as a target for new drugs or novel therapies. Our study defines a novel pathway that specifically regulates wrapping glia growth, providing a new mechanistic insight on the process by which axons in the PNS are ensheathed *in vivo*. Ral GTPase has successfully been used as a target for the design of small molecules that interfere with its function (Yan et al. 2014). Therefore, if the molecular players required for this process are conserved, *Drosophila* PNS nerves can be used as an *in vivo* system to test genetic interactions and/or new drugs and easily assess the degree of ensheathment, not only in invertebrates, but also in vertebrates.

## Materials and methods

### Fly Stocks and husbandry

All the flies used in this study were kept at 25°C, except the RNAi crosses that were performed at 29°C. The following fly strains were used: *w*^*1118*^ (BDSC 3605); *ral*^*G05010*^ (BDSC 12283); *ral*^*EE1*^ (BDSC 25095); *sec8*^*PI*^ (BDSC 12937); *sec5*^*E10*^ (Murthy et al. 2003), *sec6*^*Ex15*^ (Murthy et al. 2005); *Nrv2-GAL4* (BDSC 6800); *Repo-GAL4* (BDSC 7415); *nSyb-GAL4* (BDSC 19183); *moody-GAL4* (Stork et al. 2008); *perineurial-GAL4* (Bsg. DGRC-Kyoto 105188); Vkg-GFP (VDRC 318167); UAS-RalHA (Lee & Schwarz 2016); UAS-CD4-GFP (BDSC 35836); NrxIV-GFP (Buszczak et al. 2006), UAS-FasII-Flag (Bornstein et al. 2015). RNAi knockdown strains UAS-Ral-IR (BDSC 29580) and UAS-Sec5-IR (VDRC-*w*^*1118*^; P(GD13789) v28873).

#### Description of *Ral* mutations

*ral*^*G0501*^ is a protein null mutant (Teodoro et al. 2013) with P-element inserted upstream of the *rala* 5’ UTR region, and *Ral*^*EE1*^ harbors a point mutation that results in an amino acid substitution, Ser154Leu, within a conserved amino acid sequence predicted to be required for nucleotide binding (Eun et al. 2006), resulting in a strong reduction of the levels of this mutated protein (not shown). The *rala* gene is located on the X chromosome and the mutants analyzed in this study are either late 3^rd^ instar/early pupae lethal (*ral*^*G0501*^) or survive to adulthood where males are sterile and die a few hours post-eclosion (*ral*^*EE1*^). Because *ral* is located on the X-chromosome, we always tested male larvae. Using these two independent mutants assures that our observations are not due to 2^nd^ site mutations.

### Immunofluorescence

Larvae of the desired stage were dissected exposing the body walls, the brain and the peripheral nervous system. Dissections were performed on sylgard plates, or sylgard-covered slides, in a drop of PBS 1X and fixed in 4% formaldehyde for 20 minutes at room temperature. Antibody staining was performed in PBS containing 0.3% Triton X-100 and 5% normal goat serum. Larvae were incubated overnight at 4°C in primary antibodies, washed for at least 1h, blocked for 30min to 1h and incubated for 2h at room temperature in secondary antibodies, diluted in blocking solution. Larval fillets were mounted in mounting media DABCO (1,4-Diazabicyclo [2.2.2] octane, Sigma-Aldrich). The following primary antibodies were used: rabbit anti-HA (C29F4 Cell Signaling Technology, 1:500); mouse anti-Coracle (DSHB C615.26, 1:50); mouse anti-Futsch (DSHB 22C10, 1:100), mouse anti-Fasciclin II (DSHB 1D4, 1:100). Secondary antibodies used were the following: Alexa 488 anti-mouse and DyLight 647 anti-mouse (Jackson ImmunoResearch, 1:500). Cyanine 3 (Cy3) or Alexa 488-conjugated goat anti-HRP (Jackson ImmunoResearch, 1:500) was used to label neuronal membranes. Texas Red conjugated Phalloidin (Molecular Probes, 1:500). All reactions were performed using minimum crossreactivity secondaries. Endogenous GFP fluorescence was used in the experiments.

#### Anti HRP staining

Anti-Horseradish peroxidase (HRP) antibodies have been shown to label several neural-specific glycoproteins present in the central and peripheral nervous system of *Drosophila* and other insects (Snow et al. 1987; L. Y. Jan & Y. N. Jan 1982).

### Transmission Electron Microscopy

For TEM, larvae fillets from third instar larvae were dissected and fixed in 2% formaldehyde, 2.5% glutaraldehyde and 0.03% of picric acid in PHEM buffer at room temperature for 2h as described in Matzat *et. al*, 2015. Dissection pins were then removed and larvae were fixed overnight at 4°C with the same solution freshly made. The specimens were subsequently fixed with 2% tannic acid, 30min on ice and post-fixed with 2% Osmium reduced with 0,8% Potassium Ferricyanide during 1h on ice and 2% of uranyl acetate during 30min on ice. Larvae were dehydrated in an ascending ethanol series, were then treated with propylene and embedded in EPON resin. Transversely ultrathin sections (85-90nm) were cut 2-10 μm after the end of the ventral nerve cord. TEM images were acquired using Hitachi H-7650 microscope. For statistical analysis, we measured the area of the electron transparent regions (ETR) that is present immediately after the subperineurial glia layer (and within the wrapping glia layer) and divided this value by the total nerve area.

### Crawling/Locomotion Assay

To assess larvae behavior an open field arena crawling assay was used. Five third instar larvae were placed in a 3% agarose (MultiPurpose Agarose, tebu-bio) circular arena with a 10% agarose salt concentrated barrier on the edges of the arena. After acclimatization to the arena, larvae were filmed for 5 min at 5 frames/second. The larvae were tracked in each frame using ID Tracker software (rez-Escudero et al. 2014), which provides the coordinates used to calculate total distance and medium velocity. To represent the trajectories, the initial coordinates were plotted in a smooth line dispersion graph.

#### Locomotor Behavior Protocol - Analyses and Corrections

With the coordinates from the tracking program, medium velocity, total distance and trajectories were calculated and represented. First, the coordinates in pixels were converted in mm by applying the following formula:

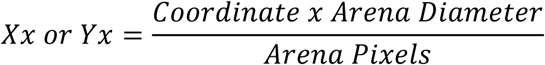

To calculate the distance, in mm, first, the distance between two points is calculates using:

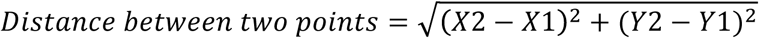

Where X and Y represent the coordinates in mm. Then, all the distances between two points are calculated and this value is the total distance of the larvae.

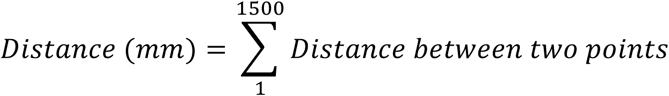

To calculate medium velocity, the distance on each two points calculated by the formula above, is divided by the time between frames. All of these values were added, forming the velocity of the larvae, as described below:

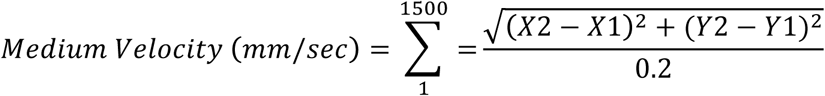

For the trajectories representation, the initial coordinates were plotted in a smooth line dispersion graph.

### Confocal imaging and quantification of nerve parameters

All images were acquired using a LSM710 confocal microscope (Zeiss). In Figure 1B, we used a 10X objective and optical sections of 10 μm; images for Supplementary Figure 2 with segmental nerves were acquired with a 40X objective with optical sections of 0.7μm. All intersegmental nerves (ISN) were acquired using a 63X lens, with optical sections of 0.7μm. All images were acquired using the same settings within each experiment. Images were analyzed using maximum z-stack projection in ImageJ.

#### Quantification of nerve thickness

nerve thickness was measured in the ISN at the region of muscle 4 innervation, segments A2-A4. We choose for measurement a region in the maximum projection image that is representative of the ISN and that is not at the motor neuron exit point and does not include any glial nuclei, since these can represent thicker points in the nerve. We used the measure tool in ImageJ to acquire the thickness (in μm).

#### Quantification of 80 μm length nerve area

we used ImageJ to trace a rectangle that was 80 μm in length and 20 μm in height to assure that the entire region of interest (ROI) is captured. We then use the threshold function followed by a *convert to binary,* to outline the entire ROI in the HRP channel (HRP ROI): this labels the whole HRP area in each genotype. We then do a measure the area using the ImageJ *measure* function, limited to the HRP ROI, to determine the total nerve area within 80 μm length.

#### Measurement of ISN FasII intensity

we drew a rectangle of 80 μm in length, similar to nerve area measurements but, because we wanted to quantify the intensity of FasII within the axons, we limited the height of the rectangle to 4 μm (this is especially important in our experiments because *ral* mutants have thicker nerves: if we divide an identical value of axonal FasII by a larger HRP ROI present in *ral* mutants, it results in artificially low values of FasII). Following the determination of the HRP ROI, using the same procedure as for the nerve area measurements, we took a FasII intensity measure within the HRP ROI.

### Statistical analysis

Statistical analysis was conducted using Excel (Microsoft Corporation, 2011) and/or GraphPad Prism 6 software (Graphpad Software, La Jolla, CA). Datasets were tested for normality and if all samples passed this test, we used parametric tests; otherwise, non-parametric tests were used. For pairwise comparisons, statistical significance was determined using a Mann-Whitney test. All multiple comparisons were performed using the following criteria: 1) if values normally distributed, a one-way analysis of variance (ANOVA) with Bonferroni multiple comparisons test, or 2) if values not normally distributed, Kruskal-Wallis with Dunn’s multiple comparisons test. Data is represented with box plots which comprise the 25–75% coincidence interval, as well as the median. Whiskers comprise the 90% coincidence interval. In the legend and text mean ± s.e.m.; sample size (n) is reported. In Figure 4C measurements are shown as mean ± s.e.m.; sample size (n) is described either in the figure legend or in Results section. All mean ± s.e.m. represented in the figure graphs are provided in Supplementary Data 1.

## Supporting information

Silva-Rodrigues et al._Supplemetal Figures

## Acknowledgements

We would like to thank César Mendes for help with the behavioral experiments and for critical discussions throughout, to AYG, Alisson Gontijo, Catarina Homem and Lara Carvalho for critical reading of the manuscript. We thank Telmo Pereira from the Microscopy Facility for technical support. We thank Marta Santos and the Fly Facility; CONGENTO: consortium for genetically tractable organisms. We thank Oren Schuldiner, Thomas Schwarz, the Developmental Studies Hybridoma Bank, the Drosophila Genomics Resource Center, and the Bloomington Drosophila Stock Center for antibodies, and fly stocks. This work was supported by H2020-GA661543-Neuronal Trafficking (ROT), IF/00392/2013/CP1192/CT0002, FCT – Portugal (ROT) and by UID/Multi/04462/2013 from iNOVA4Health (co-funded by FCT-FEDER-PT2020).

## Ethics Statement

Not Applicable.

