## Supplementary material for "Peripheral axonal ensheathment is regulated by Ral GTPase and the exocyst complex": Silva-Rodrigues et al._Supplemetal Figures

**Supplementary Figure 1**

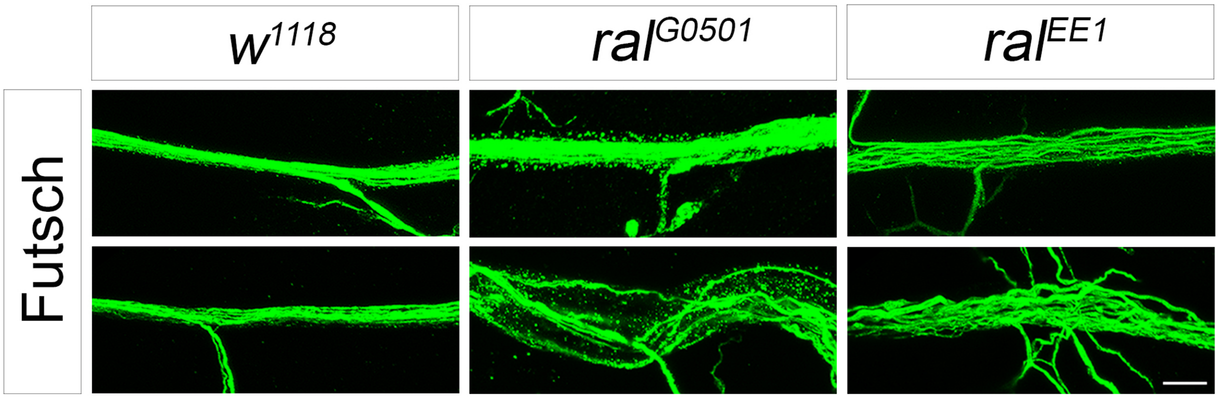

**Supplementary Figure 1. *ral* mutants exhibit some degree of axonal defasciculation.**

Axonal defasciculation is often associated with the thickening of nerve bundles. **(A)** Representative examples of different degrees of axonal defasciculation defects in the following genotypes: $w^{1118}$, $ral^{G0501}$ and $ral^{EE1}$. We used the microtubule associated protein Futsch to label axons (anti-Futsch in green). Scale bar: 10μm.

#### Supplementary Figure 2

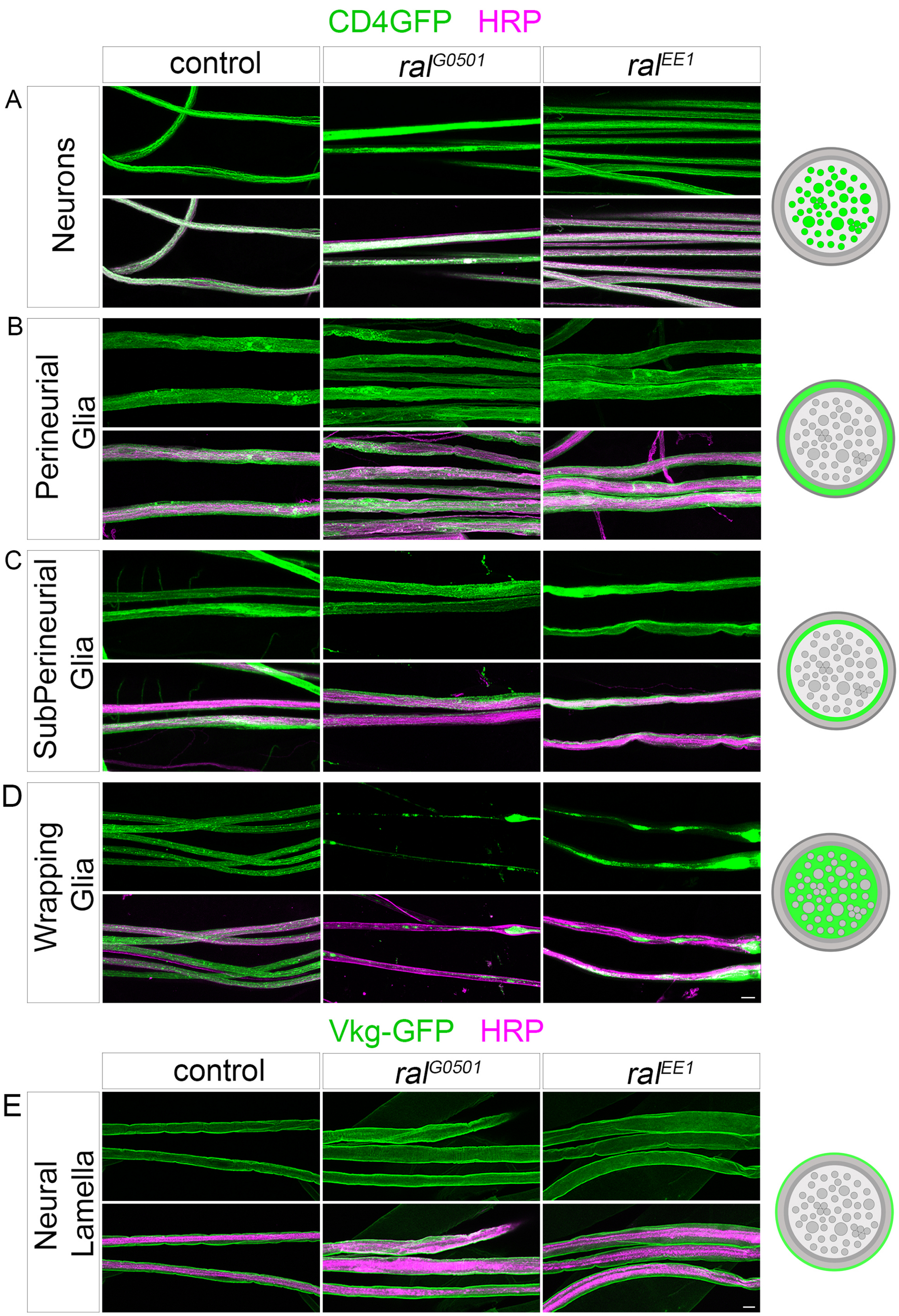

**Supplementary Figure 2. *ral* mutants have defective wrapping glia in peripheral nerves.**

**(A-D)** Analysis of the morphology of **(A)** Neurons, **(B)** Perineurial Glia, **(C)** Subperineurial Glia and **(D)** Wrapping Glia in $3^{rd}$ instar peripheral nerves. UAS-CD4-GFP was used as a membrane marker, and cell type specific Gal4 drivers (nSyb-Gal4>neurons, Bsg-Gal4> PG, moody-Gal4>SPG and Nrv2-Gal4>WG) were used. Membranes of each cell type were visualized by CD4-GFP (in green) and segmental nerves visualized with HRP (in magenta). For neural lamella **(E)** the protein trap Vkg-GFP (in green) was used. The labelling was performed in $w^{1118}$, $ral^{G0501}$ and $ral^{EE1}$ mutant background. All cells with the exception of wrapping glia are morphologically normal when comparing to their respective control. Wrapping glia are strikingly underdeveloped and fail to wrap the axons within the bundle in $ral^{G0501}$ and $ral^{EE1}$ larvae. Scale bar is 10 µm. On the right side of the figure panels, is an illustration of the cell type labeled with CD4-GFP (in green) in a cartoon of a nerve cross-section, the non-labeled layers are represented in grey.

### Supplementary Figure 3

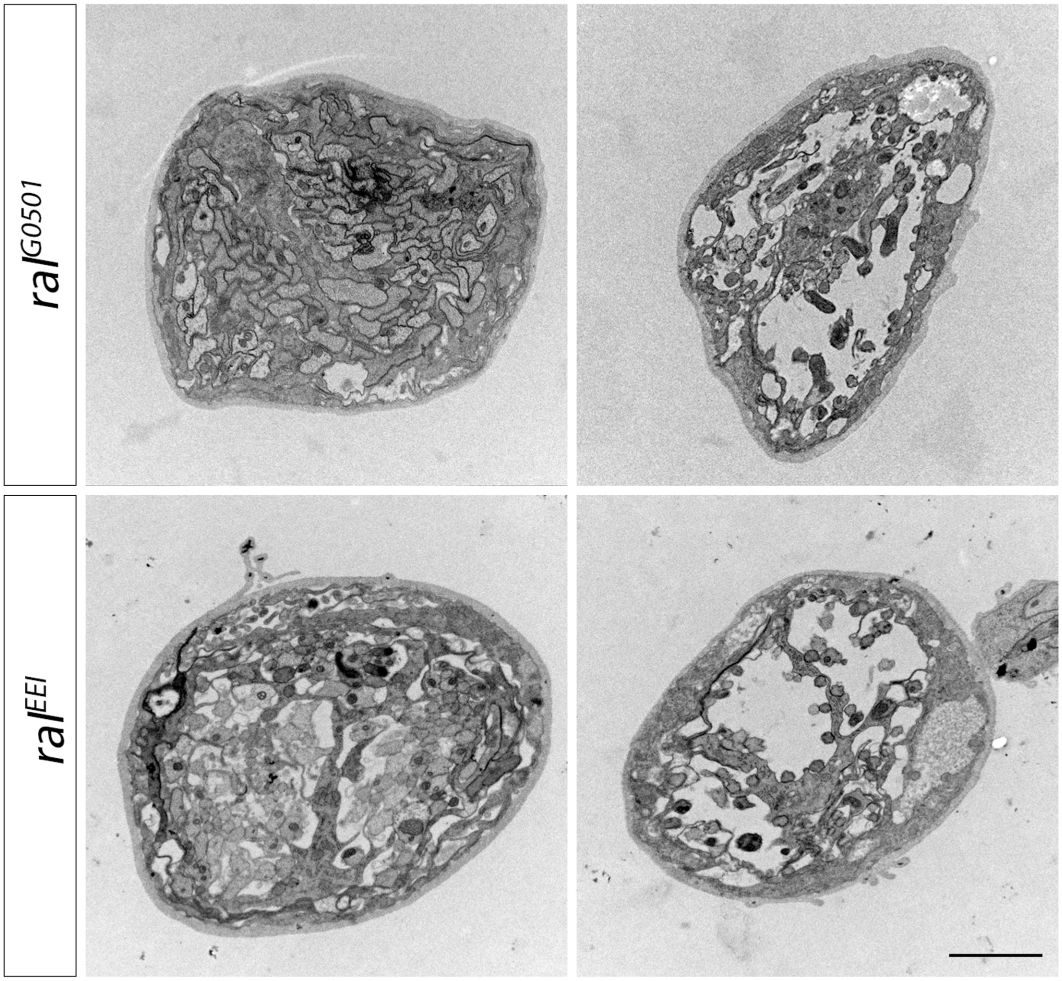

**Supplementary Figure 3. *ral* mutants show signs of a disrupted Blood-Nerve-Barrier (BNB).**

Additional representative examples of TEM images of transversal cuts of segmental nerves within 2 to 10 μm from the exit of the VNC of 3rd instar larvae. The images show examples of $ral^{G0501}$ and $ral^{EE1}$ larvae, to show the range of electron-transparent regions present in *ral* mutants, which we suggest represent a partial disruption of the BNB. Scale bar is 2 μm.

**Supplementary Figure 4**

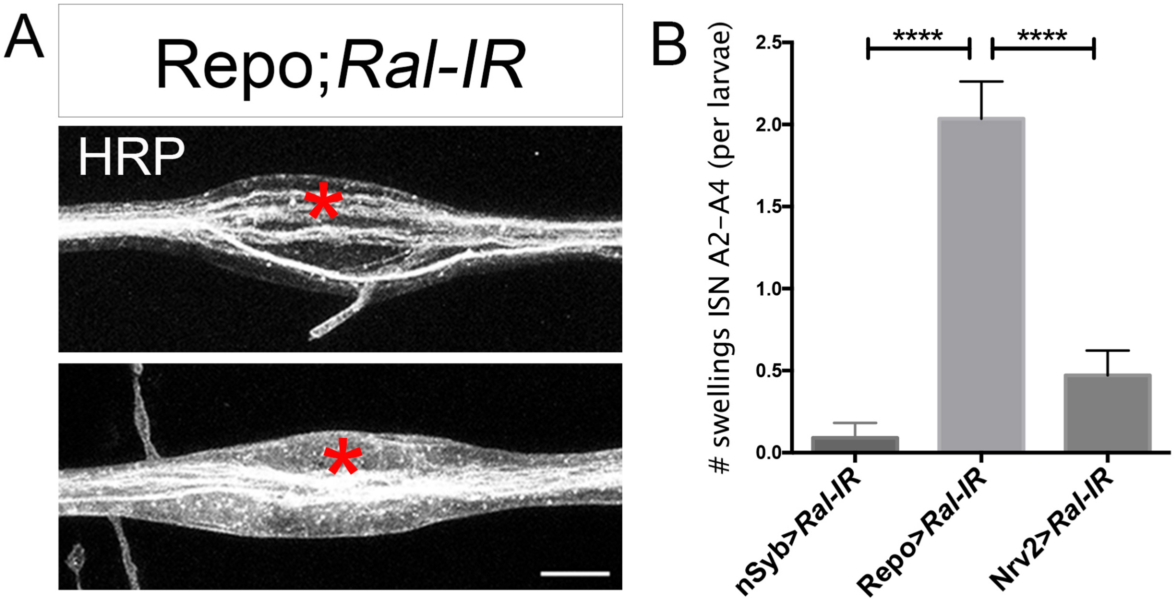

**Supplementary Figure 4. Ral reduction in all glia leads to the appearance of swellings along the ISN.**

**(A)** Examples of swellings along the ISN observed when Ral-IR is expressed in all glia (Repo-Gal4>Ral-IR), but not in control or wrapping glia Ral-IR. Hrp (grey) is used to outline the ISN. Scale bar is 10 µm. **(B)** Quantification of the number of swellings observed per larvae analyzed in the genotypes nSyb>Ral-IR (n=11), Repo>Ral-IR (n=28) and Nrv2>Ral-IR (n=17). nSyb>RalIR=0.09+0.09 / Repo>RalIR=2.04+0.23 / Nrv2>RalIR=0.47+0.79 (values in µm, mean±s.e.m.). We used a non-parametric ANOVA, Kruskal Wallis with Dunn's multiple comparisons test. ****$p<0.0001$.

#### Supplementary Figure 5

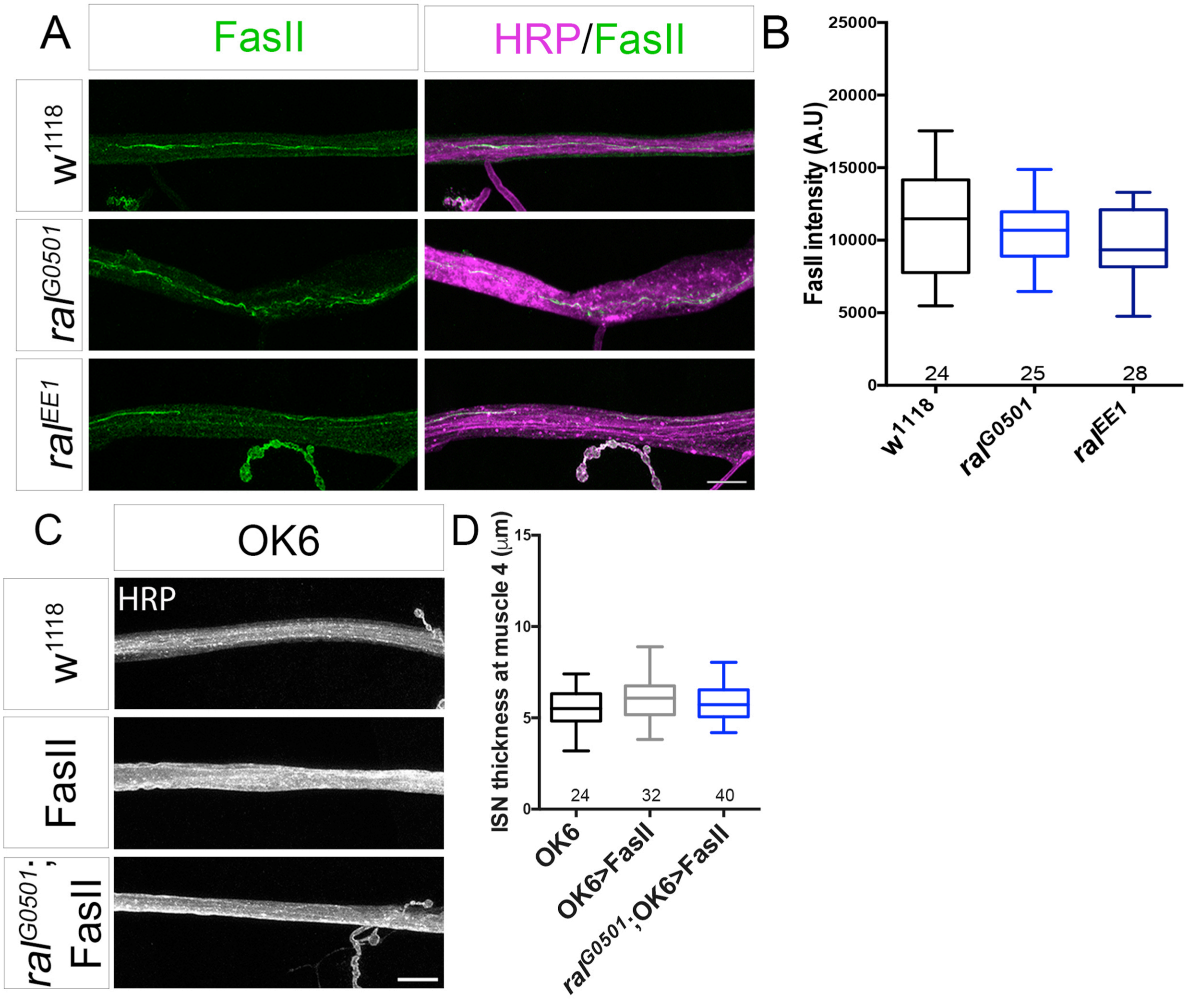

**Supplementary Figure 5. The adhesion protein Fasciclin II (FasII) levels is unchanged in *ral* mutants but its over-expression in motor neurons rescues ISN thickening.**

**(A)** Representative examples of the ISN labeled with anti-FasII antibody in wild-type ($w^{1118}$), $ral^{G0501}$ and $ral^{EE1}$ mutants. **(B)** Quantification of FasII intensity within the axonal region of the ISN at m4 exit point, shows that Fas II levels are identical in WT and *ral* mutant larvae. **(C)** Overexpression of UAS-FasII specifically in motor neurons using OK6-Gal4 to promote adhesion between motor neurons. Representative examples of the ISN labeled with HRP (grey) to outline the nerve in OK6, OK6>FasII and in $ral^{G0501}$;OK6>FasII. Scale bar is 10μm. **(D)** Quantifications of the width of the ISN at m4 exit point of the indicated genotypes. OK6=5.56±0.23, OK6>FasII=6.09±0.22, $ral^{G0501}$;OK6>FasII=5.85±0.16 (values in μm, mean±s.e.m.). This analyses shows that forcing adhesion between motor neurons is able to induce the recovery of the thick ISN present in *ral* mutants (Figure1).

**Supplementary Data 1.** Values of parameters quantified in the Figure graphs:

**Figure 1**

(D) ISN thickness

$w^{1118}$=5.21±0.08, $ral^{G0501}$=8.01±0.26 and $ral^{EE1}$=7.82±0.13

(± represent s.e.m., values in μm).

(E) ISN area in 80μm of nerve

$w^{1118}$=446.9±9.9, $ral^{G0501}$=670.0±30.9 and $ral^{EE1}$=653.4±12.9

(± represent s.e.m., values in $\mu m^2$).

**Figure 4**

(C) ETR/total nerve area

$w^{1118}$=0.93±0.33, $ral^{G0501}$=6.55±1.49 and $ral^{EE1}$=10.28±2.31

(± represent the s.e.m., values in % of total nerve area).

**Figure 5**

(B) Distance

$w^{1118}$=134.6±7.7, $ral^{G0501}$=92.5±5.7, $ral^{EE1}$=88.2±4.4, values in mm ± s.e.m.

Average speed

$w^{1118}$=0.45±1.05, $ral^{G0501}$=0.36±0.69, $ral^{EE1}$=0.37±0.67, values in mm/s ± s.e.m.

Speed of forward crawls

$w^{1118}$=0.67±0.07, $ral^{G0501}$=0.37±0.02, $ral^{EE1}$=0.38±0.04, values in mm/s ± s.e.m.

**Figure 6**

(C) ISN thickness

Ral-IR=5.13±0.11; nSyb=5.66±0.19; nSyb>Ral-IR=6.91±0.28; Repo=4.70±0.15;

Repo>Ral-IR=7.87±0.32; Nrv2=5.21±0.13; Nrv2>Ral-IR=7.13±0.24

(values in μm±s.e.m.).

(D) ISN area in 80μm of nerve

Ral-IR=381.1±9.4; nSyb=482.7±23.8; nSyb>Ral-IR=577.3±30.8; Repo=353.8±10.8;

Repo>Ral-IR=608.5±21.1; Nrv2=438.6±11.6; Nrv2>Ral-IR =549.9±15.4

(values in $\mu m^2$±s.e.m.).

**Figure 7**

(C) ISN thickness

Values for $ral^{G0501}$ graph are: RalHA=5.54±0.13, $ral^{G0501}$;RalHA=8.03±0.20, $ral^{G0501}$;RalHA>nSyb-Gal4=6.98±0.24, $ral^{G0501}$;RalHA>Repo-Gal4=6.11±0.15, $ral^{G0501}$;RalHA>Nrv2-Gal4=6.17±0.19.

Values for $ral^{EE1}$ graph are: RalHA=5.14±0.12, $ral^{EE1}$;RalHA=7.19±0.19, $ral^{EE1}$;RalHA>nSyb-Gal4=6.10±0.17, $ral^{EE1}$;RalHA>Repo-Gal4=5.85±0.18, $ral^{EE1}$;RalHA>Nrv2-Gal4=6.77 ±0.22.

(values in μm±s.e.m.).

(D) ISN area in 80μm of nerve

For $ral^{G0501}$ graph: RalHA=400.9±10.5, $ral^{G0501}$;RalHA=5.77.5±10.0, $ral^{G0501}$;RalHA>nSyb-Gal4=507.6±14.2, $ral^{G0501}$;RalHA>Repo-Gal4=460.3±12.1, $ral^{G0501}$;RalHA>Nrv2-Gal4=481.9±12.7; for $ral^{EE1}$ graph: RalHA=394.4±10.7, $ral^{EE1}$;RalHA=532.0±15.4, $ral^{EE1}$;RalHA>nSyb-Gal4=498.3±14.3, $ral^{EE1}$;RalHA>Repo-Gal4=428.2±12.0, $ral^{EE1}$;RalHA>Nrv2-Gal4=529.5±13.6

(values in $\mu m^2$±s.e.m.).

**Figure 8**

(B) ISN thickness

Control=5.66±10.26, $sec8^{P1}$=4.66±11.10

(values in μm±s.e.m.).

(D) ISN thickness

nSyb=5.92±0.25; nSyb>Sec5-IR=5.41±0.17, Repo=4.64±0.10, Repo>Sec5-IR=8.02±0.37, Nrv2=6.40±0.25, Nrv2>Sec5-IR=7.48±0.19.

(values in μm±s.e.m.).

**Figure 9**

(B) ISN thickness

$w^{1118}$=5.21±0.08, $sec5^{E10}$,$sec6^{Ex15}$/+=5.93±0.12, $ral^{G0501}$/+=5.52±0.16, $ral^{EE1}$/+=5.02±0.18, $ral^{G0501}$/+;$sec5^{E10}$,$sec6^{Ex15}$/+=7.49±0.19, $ral^{EE1}$/+;$sec5^{E10}$,$sec6^{Ex15}$/+=7.13±0.21

(mean±s.e.m., μm).
